# Mucosal Adenoviral-vectored Vaccine Boosting Durably Prevents XBB.1.16 Infection in Nonhuman Primates

**DOI:** 10.1101/2023.11.06.565765

**Authors:** Matthew Gagne, Barbara J. Flynn, Shayne F. Andrew, Dillon R. Flebbe, Anna Mychalowych, Evan Lamb, Meredith E. Davis-Gardner, Matthew R. Burnett, Leonid A. Serebryannyy, Bob C. Lin, Laurent Pessaint, John-Paul M. Todd, Zohar E. Ziff, Erin Maule, Robin Carroll, Mursal Naisan, Yogita Jethmalani, James Brett Case, Igor P. Dmitriev, Elena A. Kashentseva, Baoling Ying, Alan Dodson, Katelyn Kouneski, Nicole A. Doria-Rose, Sijy O’Dell, Sucheta Godbole, Farida Laboune, Amy R. Henry, Josue Marquez, I-Ting Teng, Lingshu Wang, Qiong Zhou, Bushra Wali, Madison Ellis, Serge Zouantchangadou, Alex Van Ry, Mark G. Lewis, Hanne Andersen, Peter D. Kwong, David T. Curiel, Kathryn E. Foulds, Martha C. Nason, Mehul S. Suthar, Mario Roederer, Michael S. Diamond, Daniel C. Douek, Robert A. Seder

**Affiliations:** Vaccine Research Center, National Institute of Allergy and Infectious Diseases, National Institutes of Health; Bethesda, MD 20892, USA; Center for Childhood Infections and Vaccines of Children’s Healthcare of Atlanta, Department of Pediatrics, Emory University School of Medicine; Atlanta, GA 30322, USA; Emory Vaccine Center, Emory University; Atlanta, GA 30329, USA; Emory National Primate Research Center; Atlanta, GA 30329, USA; Bioqual, Inc.; Rockville, MD 20850, USA; Department of Medicine, Washington University School of Medicine; St. Louis, MO 63110, USA; Department of Radiation Oncology, Washington University School of Medicine; St. Louis, MO 63110, USA; Biostatistics Research Branch, Division of Clinical Research, National Institute of Allergy and Infectious Diseases, National Institutes of Health; Bethesda, MD 20892, USA; Department of Pathology & Immunology, Washington University School of Medicine; St. Louis, MO 63110, USA; Department of Molecular Microbiology, Washington University School of Medicine; St. Louis, MO 63110, USA; The Andrew M. and Jane M. Bursky Center for Human Immunology & Immunotherapy Programs, Washington University School of Medicine; St. Louis, MO 63110, USA; Center for Vaccines & Immunity to Microbial Pathogens, Washington University School of Medicine; St. Louis, MO 63110, USA

**Keywords:** SARS-CoV-2, COVID-19, adenoviral-vectored vaccine, intranasal vaccine, aerosolized vaccine, mucosal immunity, IgA, secretory antibody, immune imprinting, XBB.1.16

## Abstract

Waning immunity and continued virus evolution have limited the durability of protection from symptomatic infection mediated by intramuscularly (IM)-delivered mRNA vaccines against COVID-19 although protection from severe disease remains high. Mucosal vaccination has been proposed as a strategy to increase protection at the site of SARS-CoV-2 infection by enhancing airway immunity, potentially reducing rates of infection and transmission. Here, we compared protection against XBB.1.16 virus challenge 5 months following IM or mucosal boosting in non-human primates (NHP) that had previously received a two-dose mRNA-1273 primary vaccine regimen. The mucosal boost was composed of a bivalent chimpanzee adenoviral-vectored vaccine encoding for both SARS-CoV-2 WA1 and BA.5 spike proteins (ChAd-SARS-CoV-2-S) and delivered either by an intranasal mist or an inhaled aerosol. An additional group of animals was boosted by the IM route with bivalent WA1/BA.5 spike-matched mRNA (mRNA-1273.222) as a benchmark control. NHP were challenged in the upper and lower airways 18 weeks after boosting with XBB.1.16, a heterologous Omicron lineage strain. Cohorts boosted with ChAd-SARS-CoV-2-S by an aerosolized or intranasal route had low to undetectable virus replication as assessed by levels of subgenomic SARS-CoV-2 RNA in the lungs and nose, respectively. In contrast, animals that received the mRNA-1273.222 boost by the IM route showed minimal protection against virus replication in the upper airway but substantial reduction of virus RNA levels in the lower airway. Immune analysis showed that the mucosal vaccines elicited more durable antibody and T cell responses than the IM vaccine. Protection elicited by the aerosolized vaccine was associated with mucosal IgG and IgA responses, whereas protection elicited by intranasal delivery was mediated primarily by mucosal IgA. Thus, durable immunity and effective protection against a highly transmissible heterologous variant in both the upper and lower airways can be achieved by mucosal delivery of a virus-vectored vaccine. Our study provides a template for the development of mucosal vaccines that limit infection and transmission against respiratory pathogens.

**Graphical abstract:** 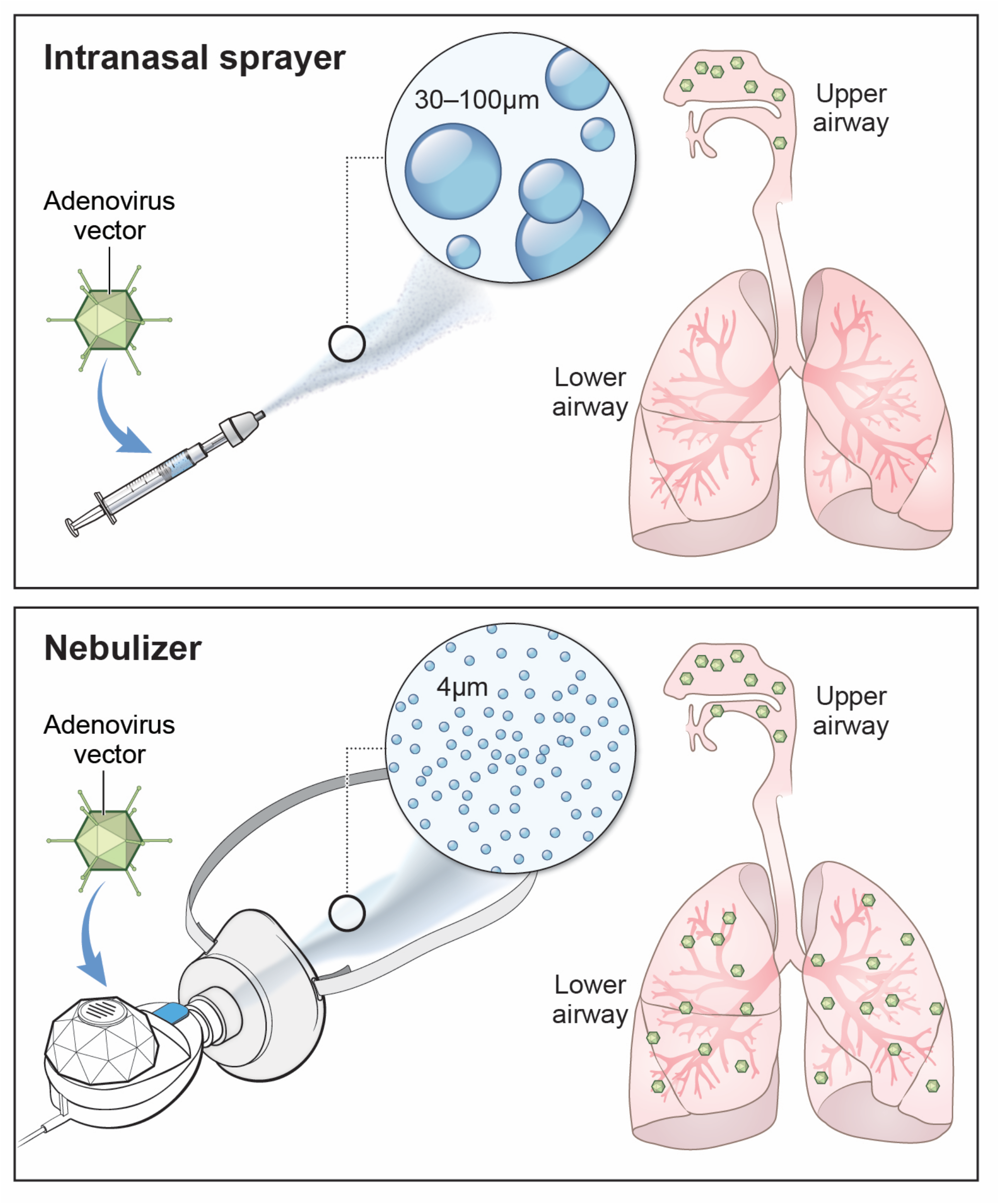

## Introduction

Immunity to SARS-CoV-2 elicited by the majority of approved vaccine boosts given by the intramuscular (IM) route continues to protect against severe disease and hospitalization^1, 2, 3, 4^, which remain important goals of vaccination against SARS-CoV-2. However, breakthrough infections occur frequently, which has sustained the COVID-19 pandemic. Indeed, effectiveness of bivalent mRNA vaccines against infection approaches 0% within four months of immunization^5^. Furthermore, re-infections are associated with post-acute sequelae, often termed long COVID^6^. In addition, the elderly and immunocompromised remain susceptible to severe disease due to poor vaccine responses or diminished innate immunity^7, 8, 9, 10^. Thus, deployment of vaccines that prevent or substantially limit breakthrough infections and subsequent transmission could be clinically beneficial, slow the emergence of new strains and alter the dynamics of population-wide virus spread.

Such a vaccine would need to overcome four major obstacles contributing to breakthrough infection: (1) waning of neutralizing antibody titers with time^11, 12, 13, 14^; (2) requirement of a high level of serum neutralizing antibodies for protection in the nose as compared to the lungs^15, 16^; (3) sustained SARS-CoV-2 evolution with the continual emergence of antigenically mismatched variants that exhibit immune escape^17, 18, 19, 20, 21^; and (4) antigenic imprinting, which could limit the induction of new variant-specific neutralizing antibody responses^22, 23, 24^.

The initial finding that a higher titer of serum antibodies is required to confer protection in the upper airway than in the lower airway^15, 16^ suggested potentially different mechanisms of virus control in distinct tissue compartments. Indeed, neutralizing antibodies, Fc effector function and T cells all may contribute to protection in the lung against severe disease^15, 25, 26, 27, 28^. By contrast, their respective contributions to upper airway protection remain less clear.

Targeting the upper or lower airways directly with a vaccine could enhance immune responses to SARS-CoV-2 by eliciting or boosting central and effector memory T cells, resident memory B cells and plasma cells in the mucosa-associated lymphoid tissue (MALT) and draining lymph nodes, and by locally generating inhibitory IgA^29, 30, 31, 32^. Many intranasal (IN) or aerosolized (AE) vaccines are under investigation including soluble spike (S)^33^ and lipid nanoparticle-encapsulated mRNAs^34^. Replication-competent or incompetent virus-vectored vaccines are an alternative approach, as these platforms can elicit local humoral and cellular immunity at the site of respiratory virus infection. Current candidates include parainfluenza virus^35^, Newcastle disease virus^36^ and numerous adenovirus serotypes^37, 38^. The vaccine ChAd-SARS-CoV-2-S contains the prefusion-stabilized S from the ancestral Wuhan strain inserted into a replication-deficient subgroup E Simian-Ad36 adenovirus and has been approved for use in India as an IN vaccine or booster administered as nasal drops^39^. ChAd-SARS-CoV-2-S elicits mucosal and systemic S-specific IgA, IgG and T cell responses and durably protected against multiple SARS-CoV-2 variants in both the upper and lower airways in preclinical animal models^40, 41, 42, 43, 44^.

NHP models have been important for the development of SARS-CoV-2 vaccines and have largely predicted immunity and protection seen in humans although disease severity is more limited^45^. NHP have been used to demonstrate vaccine-mediated protection against virus variants, define B cell imprinting and provide evidence for the importance of boosting for protection ^26, 46, 47, 48, 49, 50, 51, 52^. However, there have been few studies in NHP assessing how mucosal vaccines influence immunity and protection or their effects as a booster following a standard mRNA vaccine regimen similar to that used in humans. To address the protective role of local humoral and T cell immunity arising from a mucosal boost and determine the optimal mucosal delivery route and device, we administered a bivalent Wuhan-1 and Omicron BA.5-matched ChAd-SARS-CoV-2-S vaccine to rhesus macaques ~7 months after they had been primed with two IM doses of ancestral S-matched mRNA-1273 vaccine. One group of animals received ChAd-SARS-CoV-2-S delivered by a clinically approved aerosol device to the upper and lower airways, whereas another received the vaccine as a mist via the IN route by a clinically approved intranasal spray. For comparison, a cohort of animals were boosted via IM route with a matched bivalent mRNA vaccine (mRNA-1273.222). Four and a half months later, all animals were challenged with the highly transmissible, heterologous XBB.1.16 strain. The data presented here show that mucosal boosting confers durable and highly effective protection with the potential to block both infection and transmission.

## Results

### Study design

20 Indian-origin rhesus macaques were immunized with 30 µg of Wuhan-1/WA1 S-encoding mRNA-1273 at weeks 0 and 4 via IM route. The dose and regimen were chosen to approximate the immune responses elicited in humans by the standard mRNA-1273 primary series. Seven months later, the 20 NHP were separated into 3 groups (**Fig. 1**). One cohort (**IM boost**) of 8 NHP received 30 µg of bivalent Wuhan-1 and BA.5 S-encoding mRNA-1273.222 (both S-2P) via IM route to model the effect of administering a bivalent booster to immunologically imprinted vaccinees^24, 47, 53^. Another cohort (**IN boost**) of 6 NHP received a total of 10^11^ virus particles (vp) composed of an equal mixture of ChAd-SARS-CoV-2-Wuhan-1-S (S-2P) and ChAd-SARS-CoV-2-BA.5-S (S-6P, furin cleavage site mutation)^54^ delivered via intranasal sprayer using the MAD Nasal^TM^ Intranasal Mucosal Atomization Device. The MAD is designed to deliver particles of 30-100 µm to the upper airway as a mist. The final 6 primed NHP (**AE boost**) were boosted with the same dose of the bivalent ChAd-SARS-CoV-2-S cocktail as aerosolized 4 µm particles via an Investigational eFlow Nebulizer System (PARI Respiratory Equipment, Inc. (PARI)) with a silicone face mask attachment (PARI SMARTMASK Baby/Kids) to enable particle deposition into the nose and lungs. To interrogate the impact of mRNA vaccine-priming and the potential for use of a mucosally-delivered virus-vectored vaccine in an unexposed population, we also administered a single dose of the same bivalent ChAd-SARS-CoV-2-S vaccine to 4 naïve NHP via aerosol to the nose and lungs (**AE prime**). As control groups, 8 naïve NHP (**control**) received two doses of 30 µg control mRNA at weeks 0 and 4. At the time of boosting, 4 of these NHP received no vaccines, whereas the other 4 were given 10^11^ vp of a control adenoviral-vectored vaccine (ChAd-Control) by the AE route.

**Figure 1.**
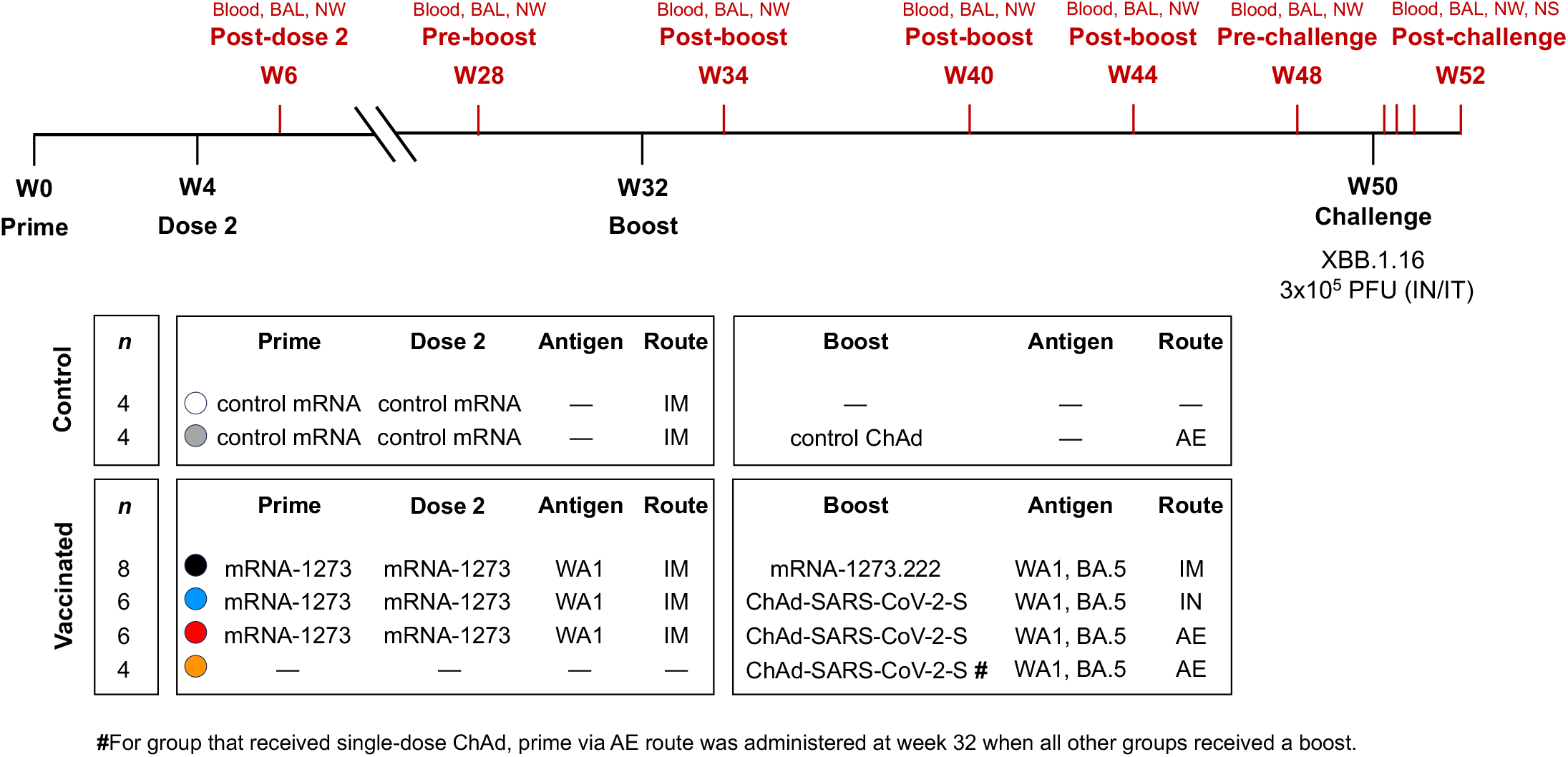
Experimental schema. NHP (*n*=8 for control groups and *n*=24 for vaccine groups) were administered 30 µg of mRNA vaccine via IM route or 1×10^11^ virus particles of adenoviral-vectored vaccine via IN or AE route according to immunization schedule shown above. Eighteen weeks after boosting, all primates were challenged with XBB.1.16. Sampling schedule indicated in red.

Samples from multiple anatomical sites, including bronchoalveolar lavage fluid (BAL), nasal washes (NW) and peripheral blood were collected for humoral and cellular analysis at week 6 (post-prime), week 28 (pre-boost), week 34 (post-boost), week 40 and 44 (memory) and week 48 (pre-challenge). At week 50, which was 4.5 months after the boost, all NHP were challenged with 3 × 10^5^ plaque forming units (PFU) of the antigen-shifted XBB.1.16 Omicron strain (**Extended Data Fig. 1**). Challenge occurred via IN and intratracheal (IT) route. BAL, NW and nasal swabs (NS) were collected post-challenge to measure levels of virus replication.

### Mucosally-delivered ChAd-SARS-CoV-2-S vaccine limits infection at the site of boosting

Subgenomic RNA (sgRNA) encoding the *N* gene is a highly sensitive and specific measure of virus replication and has been used in many prior NHP studies as the standard benchmark for assessing protection^26, 46, 55^. In the lower airway (**Fig. 2a**), sgRNA levels were similar (1.1 × 10^5^ geometric mean copies numbers per mL) between both control cohorts, which were combined for statistical analysis. On day 2, both the IM and IN boosts yielded significant protection in the lungs, with copy numbers of 9.8 × 10^2^ and 9.5 × 10^2^, respectively, which were reductions of ~115-fold compared to the controls. The AE boost group showed even greater control of virus replication, with geometric mean titers of 1.0 × 10^2^ (1,100-fold reduction), which was only marginally higher than the assay limit of detection (LOD). By day 7, 6 of 6 NHP in the AE boost group, 4 of 4 in the AE prime group, 4 of 6 in the IN boost group, and 5 of 8 in the IM boost group had cleared sgRNA from the lungs compared to only 2 of 8 animals in the control group. However, the difference in sgRNA on day 7 was statistically significant (*p* = 0.0247) only for the AE boost cohort. In summary, all vaccines tested conferred protection in the lower airways as defined by a reduction in virus replication compared to the control groups by day 2 post-challenge, consistent with clinical data on the efficacy of IM vaccines against severe disease^1, 3, 4, 56^; however, the AE boost resulted in more rapid and nearly complete control of sgRNA production.

**Figure 2.**
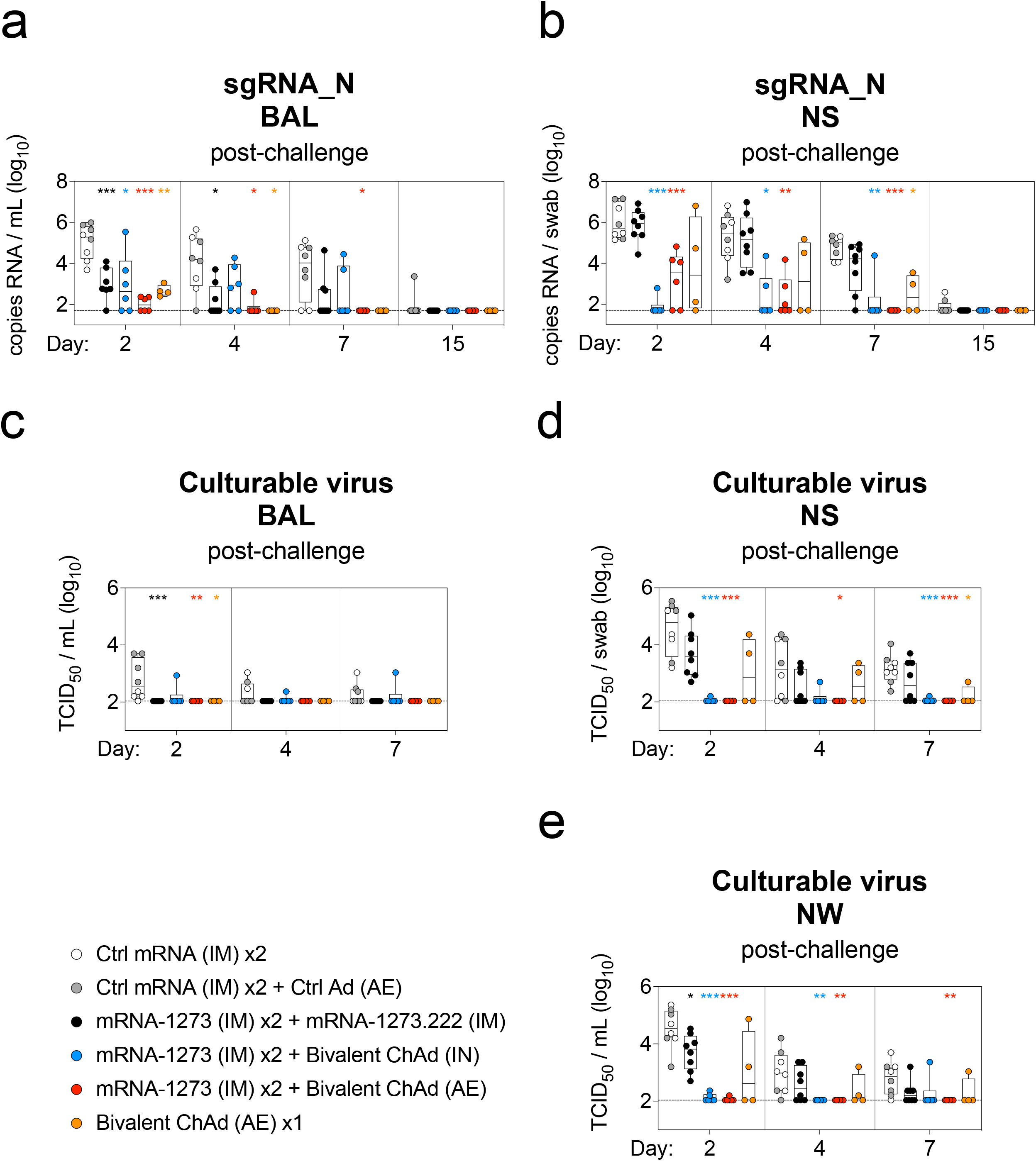
Mucosal adenoviral-vectored vaccine protects against XBB.1.16 replication. NHP (*n*=4-8 per group) were administered mRNA-1273 or control mRNA at weeks 0 and 4 and boosted at week 32 with the indicated vaccine. (**a-b**) Virus replication was measured by sgRNA_*N* assay, and (**c-e**) culturable virus was assessed by TCID_50_ assay in lower (**a, c**) and upper (**b, d, e**) airways at days 2, 4, 7 and 15 (for sgRNA only). Circles indicate individual NHP. Boxes represent interquartile range with median indicated by solid line. Assay LOD represented as dotted line. Wilcoxon Rank-Sum tests conducted for each vaccinated group in comparison to pooled controls at indicated timepoints. Additional details on statistical analysis listed in Methods. Asterisks indicate pairwise *p*-values as follows: \**p*<0.05, \*\**p*<0.01, \*\*\**p*<0.005. All other comparisons were not significant (*p*>0.05).

As the pandemic has progressed, the evolution of virus variants that resist antibody neutralization has resulted in less durable protection against symptomatic upper airway infection especially from Omicron sublineages^2, 57^. In the upper airway, geometric mean virus titers in the NS on day 2 post-challenge were 7.3 × 10^5^ in the IM boosted group and 1.0 × 10^6^ in the controls, a difference of less than 1-log_10_ (*p* > 0.05); all 8 animals in the IM boost group had detectable sgRNA in the NS (**Fig. 2b**). In contrast, there was a near-complete reduction (14,000-fold) of sgRNA in the IN boosted group with geometric mean titers of 7.6 × 10^1^ and measurable in only 1 of 6 IN-boosted NHP. sgRNA copy numbers in the NS of the AE boost group were also significantly lower at 1.8 × 10^3^ (580-fold). Even in the AE prime group with only 4 animals, there was evidence of protection, although the effect was not uniform or significant (*p* > 0.05 vs control or IM boost). By day 7 post-challenge, essentially all animals in the IN and AE boosted groups had undetectable sgRNA in the NS. By contrast, sgRNA copy numbers in the IM boosted group were reduced only 10-fold compared to the controls (*p* > 0.05). By day 15 post-challenge, sgRNA from the upper or lower airways was only detectable in primates from the control group. These data show that even ~5 months after mucosal boosting, heterologous XBB.1.16 virus replication can be effectively controlled.

To extend the analysis, we also assessed the presence of infectious virus using a tissue culture infectious dose (TCID)_50_ assay. Similar to the sgRNA findings, all mucosally vaccinated groups (boosted or primed) demonstrated similar protection in the lungs as the IM boosted group (*p* > 0.05 vs IM boost on all sampling days) (**Fig. 2c**). While there was virus detected in the BAL of 7/8 control animals, only 1 NHP across all of the vaccinated cohorts had culturable virus. In contrast, significant protection in the upper airway was observed only by the AE and IN boost groups (**Fig. 2d**). Virus titers in the NS on day 2 were 3.6 × 10^4^ in the control group compared to 1.1 × 10^2^ in the AE boosted group (330-fold reduction) and 1.2 × 10^2^ in the IN boosted group (310-fold reduction). Of note, control of virus replication was not observed in the IM boosted group on day 2. Similar findings were obtained in the NW samples (**Fig. 2e**).

### ChAd-SARS-CoV-2-S mucosally-delivered vaccine elicits durable serum humoral responses

Multiple mechanisms may account for the protective role of antibodies against SARS-CoV-2 including neutralizing and Fc effector function activities^15, 25, 26, 58, 59, 60^. Thus, we performed an extensive characterization of humoral responses in blood and mucosal tissue throughout the course of the study.

Since all boosted groups received the same 2-dose IM priming pre-boost, we present all pre-boost data as the average of all vaccinated groups, although the individual pre-boost titers are displayed in the figures for reference. Virus neutralization was measured using lentiviruses pseudotyped with S from D614G (the benchmark strain), BA.5 or XBB.1.16. Geometric mean serum neutralizing antibody titers (GMT) to D614G of 1.4 × 10^3^ reciprocal median infectious dose (ID_50_) at week 6 following IM priming decreased 6.5-fold to 2.2 × 10^2^ at week 28 (**Fig. 3a**). These data are consistent with results from clinical and pre-clinical studies in which antibody titers wane following an mRNA IM prime or boost, resulting in reduced efficacy against symptomatic infection^26, 61, 62^. Following a third IM dose (boost), serum titers were 4.5 × 10^3^ (20-fold increase). Titers to both BA.5 and XBB.1.16 were low to undetectable until the third dose, after which they reached 6.8 × 10^2^ and 1.1 ×10^2^, respectively (**Fig. 3b-c**). Two weeks after an AE boost or IN boost, serum neutralizing titers to D614G were 1.8 × 10^3^ and 9.6 × 10^2^, respectively, which were 2.6 and 4.7-fold lower than the IM boosted cohort (**Fig. 3a**). However, neutralizing titers in animals that received either of the mucosal ChAd-SARS-CoV-2-S boosts remained stable over 5 months compared to the decline following an IM boost. Indeed, at week 48, titers to D614G in the IM, IN and AE boost groups were 8.7 × 10^2^, 5.8 × 10^2^ and 1.5 × 10^3^, respectively, and to XBB.1.16 were 3.8 × 10^1^, 2.2 × 10^1^ and 5.8 × 10^1^. Neutralizing responses to emerging SARS-CoV-2 strains BA.2.86 and EG.5.1 also were assessed (**Extended Data Fig. 2**). Only the IM and AE boosted groups had detectable neutralizing titers to these variants. Two weeks after the boost, titers were similar between the groups with GMT to BA.2.86 of 7.0 × 10^1^ for the IM boost group and 4.5x 10^1^ for the AE boost group, whereas GMT to EG.5.1 were 5.7 × 10^1^ and 4.8 × 10^1^ for the same groups, respectively.

**Figure 3.**
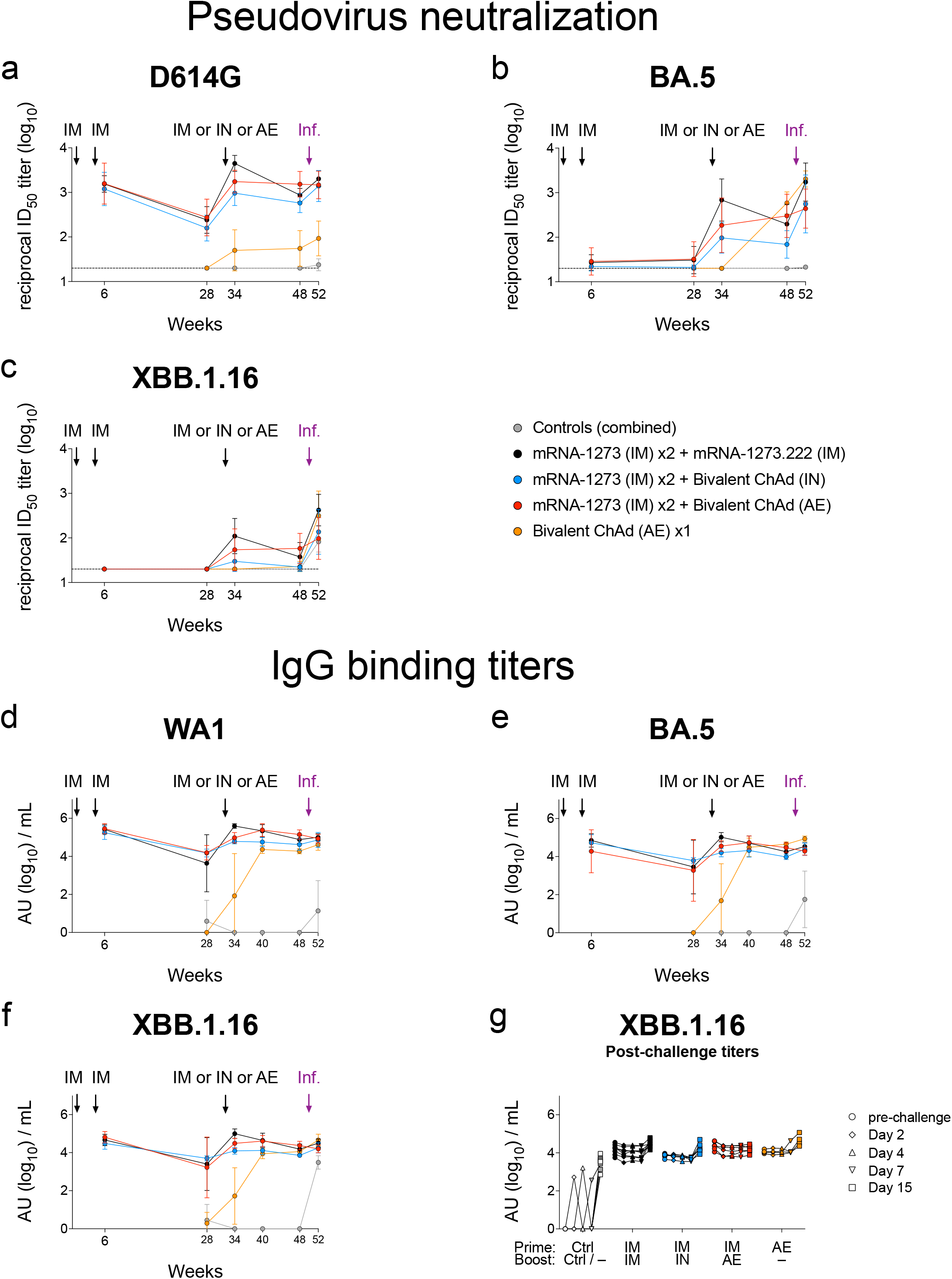
Mucosal adenoviral-vectored vaccine elicits durable systemic humoral responses. NHP (*n*=4-8 per group) were administered mRNA-1273 or control mRNA at weeks 0 and 4 and boosted at week 32 with the indicated vaccine. (**a-g**) Sera were collected post-prime (week 6), pre-boost (week 28), post-boost (week 34 and week 40), pre-challenge (week 48) and post-challenge (days 2, 4, 7 and 15). (**a-c**) Pseudovirus neutralizing responses measured against (**a**) D614G, (**b**) BA.5 and (**c**) XBB.1.16. Circles indicate geometric means for each group. Error bars represent geometric standard deviation. Assay LOD represented as dotted line. (**d-f**) Serum IgG binding titers to (**d**) WA1, (**e**) BA.5 and (**f**) XBB.1.16 S at indicated times. Circles indicate geometric means for each group. Error bars represent geometric standard deviation and may extend beyond range of graph. (**g**) Post-challenge binding titers to XBB.1.16 S for individual NHP. Pre-challenge samples collected at week 48. Symbols in **g** indicate AU / mL of individual NHP at indicated times. AU below a value of 1 were replaced with a value of 1 for data shown in **d-g**.

The AE prime group also exhibited different neutralizing titer kinetics than those observed following the IM mRNA primary series of two immunizations. Two weeks after the AE prime, neutralizing titers to D614G and BA.5 were 5.0 × 10^1^ and 2.0 × 10^1^, respectively (**Fig. 3a,b**). Fourteen weeks later, just before the time of challenge (TOC), titers to D614G were still at 5.5 × 10^1^, whereas titers to BA.5 had increased 29-fold to 5.9 × 10^2^ indicating that the peak response to the AE vaccine was considerably later than the two-week interval typically observed following IM vaccination. Indeed, at week 48, neutralizing titers to BA.5 were higher in the AE prime group than in all other cohorts, likely reflecting the lack of imprinting to ancestral epitopes from prior mRNA IM vaccination. Overall, these data are consistent with the findings in human studies that serum neutralizing titers following IM-delivered Ad26-vectored vaccines remain stable over time^63, 64^.

Analysis of serum neutralizing titers at 2 weeks post-XBB.1.16 challenge showed 11.2-fold, 6.2-fold and 13.8-fold increases in the IM boost, IN boost and AE prime vaccinated animals respectively, whereas titers in the AE boost group increased by only 1.7-fold compared to the time of challenge (**Fig. 3c**). A kinetic analysis of IgG binding titers for all vaccine groups showed similar findings to the neutralizing responses, both pre- and post-challenge (**Fig. 3d-f**). In control NHP, primary serum binding IgG responses were detected to all variant S proteins at 2 weeks post-challenge (**Fig. 3g**, **Extended Data Fig 3**), with the highest response against XBB.1.16, the challenge strain. In comparison, we detected a recall response to XBB.1.16 at two weeks post-challenge in all vaccinated groups except for the AE boost group, which showed no boost in serum anti-S IgG titers (**Fig. 3g**). These data suggest that rapid control of virus replication in the AE-boosted animals limited antigen generation and consequently, systemic boosting of neutralizing and binding antibodies.

### Analysis of upper and lower airway mucosal antibody responses

mRNA vaccines administered by the IM route either as primary series or following a boost principally elicit S-specific serum and mucosal IgG with low to undetectable levels of mucosal IgA^65, 66^. Indeed, anti-S IgG responses were boosted in the BAL following a third IM dose of mRNA, with titers to WA1 increasing from 1 × 10^0^ at week 28 to 3.8 × 10^2^ arbitrary binding units (AU) / mL at week 34 (2 weeks post-boost), which was a 380-fold increase (**Fig. 4a**). BAL IgG titers to BA.5 and XBB.1.16 also increased substantially (**Fig. 4b-c**). However, IgA titers in the lungs against WA1 S after IM boost were low and increased only ~3-fold from 1.1 × 10^0^ at week 28 to 3.6 × 10^0^ at week 34 (**Fig. 4d**). IgA titers to all other variants were also low to undetectable (**Fig. 4e-f**). In contrast, the AE boost strikingly increased both IgG (360-fold increase) and IgA titers (1,200-fold increase) against WA1 S in the BAL, and these titers were stable over the following 5 months. In the IN boost group, there were modest increases of WA1 S-specific antibody titers in BAL with fold increases of 90-fold for IgG and 26-fold for IgA compared to the pre-boost timepoint. Lastly, two weeks after the AE prime (week 34), we also detected WA1 S-specific IgG (1.9 × 10^1^) and IgA (1.6 × 10^2^) in the BAL, and these continued to rise by week 40, similar to the trend observed for serum neutralizing antibody titers. Despite having only received a single-dose of ChAd-SARS-CoV-2-S, the AE prime group had the second-highest anti-WA1 S IgG and IgA titers in the lungs at the time of challenge (**Fig. 4a,d**) and the highest anti-BA.5 S IgG titers (**Fig. 4b**).

**Figure 4.**
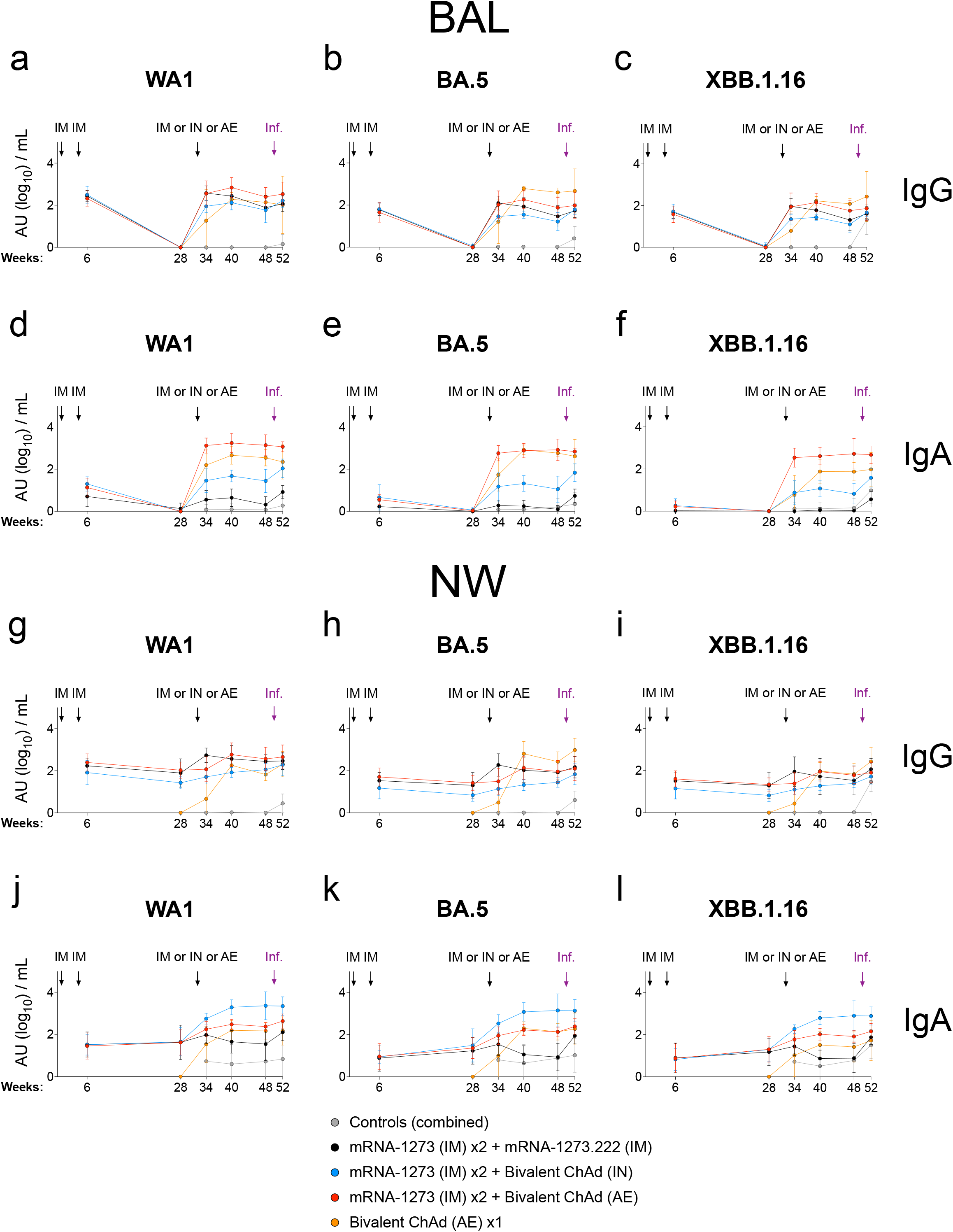
Mucosal IgG and IgA responses following vaccination. NHP (n=4-8 per group) were administered mRNA-1273 or control mRNA at weeks 0 and 4 and boosted at week 32 with the indicated vaccine. (**a-f**) BAL and (**g-l**) NW were collected post-prime (week 6), pre-boost (week 28), post-boost (week 34 and week 40), pre-challenge (week 48) and two weeks post-challenge (week 52). (**a-c, g-i**) IgG and (**d-f, j-l**) IgA binding titers measured using WA1, BA.5 or XBB.1.16 S as indicated. Circles indicate geometric means for each group. Error bars represent geometric standard deviation and may extend beyond range of graph. AU below a value of 1 were replaced with a value of 1.

In the NW, each of the boosts resulted in only modest increases in anti-S IgG binding titers (**Fig. 4g-i**), in contrast to the large increases observed in the BAL. The AE prime, however, induced upper airway WA1 S-specific IgG that reached titers equivalent to the mucosally-boosted groups by week 40. Importantly, the IN boost, but not the AE or IM boosts, increased anti-S IgA levels in the nose by week 34 with 13-fold, 15-fold and 10-fold increases against WA1, BA.5 and XBB.1.16, respectively (**Fig. 4j-l**). Anti-S IgA titers remained stable in the AE and IN boost groups until challenge.

### Kinetics of airway mucosal antibody responses post-XBB.1.16 challenge

In prior NHP studies, we have shown that there is a rapid anamnestic antibody response in BAL fluid within 2-4 days of SARS-CoV-2 challenge of IM-vaccinated animals^26, 67^ and this coincides with control of virus replication. Here, there was an increase in anti-S IgG antibodies in BAL post-challenge across all the vaccine groups (**Extended Data Fig. 4a-c**). By day 4 post-challenge, geometric mean IgG binding titers in the BAL fluid to XBB.1.16 had increased 3- to 5-fold for all vaccinated groups compared to the pre-challenge timepoints (**Extended Data Fig. 4c**), although this response was only transient in the AE boost group. Furthermore, anamnestic IgA responses in BAL fluid to any variant S were only clearly detected in the IM and IN boosted cohorts, with delayed kinetics compared to IgG (**Extended Data Fig. 4d-f**). Anti-XBB.1.16 S IgA binding titers increased 3.4-fold in the IM boost group and 5.8-fold in the IN boost group by day 15 post-challenge compared to the pre-challenge timepoint (**Extended Data Fig. 4f**). Despite the anamnestic IgA responses in the lower airway in the IN boosted group, there was no corresponding recall responses in the nose. Indeed, NW IgA was boosted by XBB.1.16 challenge only in the IM boosted group by day 15 (10-fold) (**Extended Data Fig. 4l**). This is consistent with persistent virus replication in the nose observed only in the IM boosted group, which likely provided sufficient S antigen to boost local IgA responses.

### Differential protection by mucosal IgG and IgA against SARS-CoV-2 in the upper and lower airway

To determine the potential contributions of IgG and IgA to protection in the respiratory tract, we measured the ability of mucosal fluid (BAL or NW) antibodies to inhibit binding between variant S and its angiotensin-converting enzyme 2 (ACE2) receptor as a functional surrogate of neutralization^68^. The binding inhibition assay was used as a sample-sparing measure compared to the pseudovirus neutralization assay. ACE2 binding inhibition activity in lung and nasal fluids at week 28 (pre-boost) was low to undetectable for all groups following two IM doses of mRNA-1273 (**Fig. 5**). In the IM boost group, inhibition of WA1 binding to ACE2 in the BAL fluid increased two weeks after the boost, with median binding inhibition rising from 6.9% to 58.8% (**Fig. 5a**). Binding inhibition then waned, falling to a median value of 17.6% just before challenge (week 48), consistent with the decrease in serum neutralization and binding titers. We did not observe a clear increase in inhibition of any other SARS-CoV-2 variant after IM boosting (**Fig. 5a-c**).

**Figure 5.**
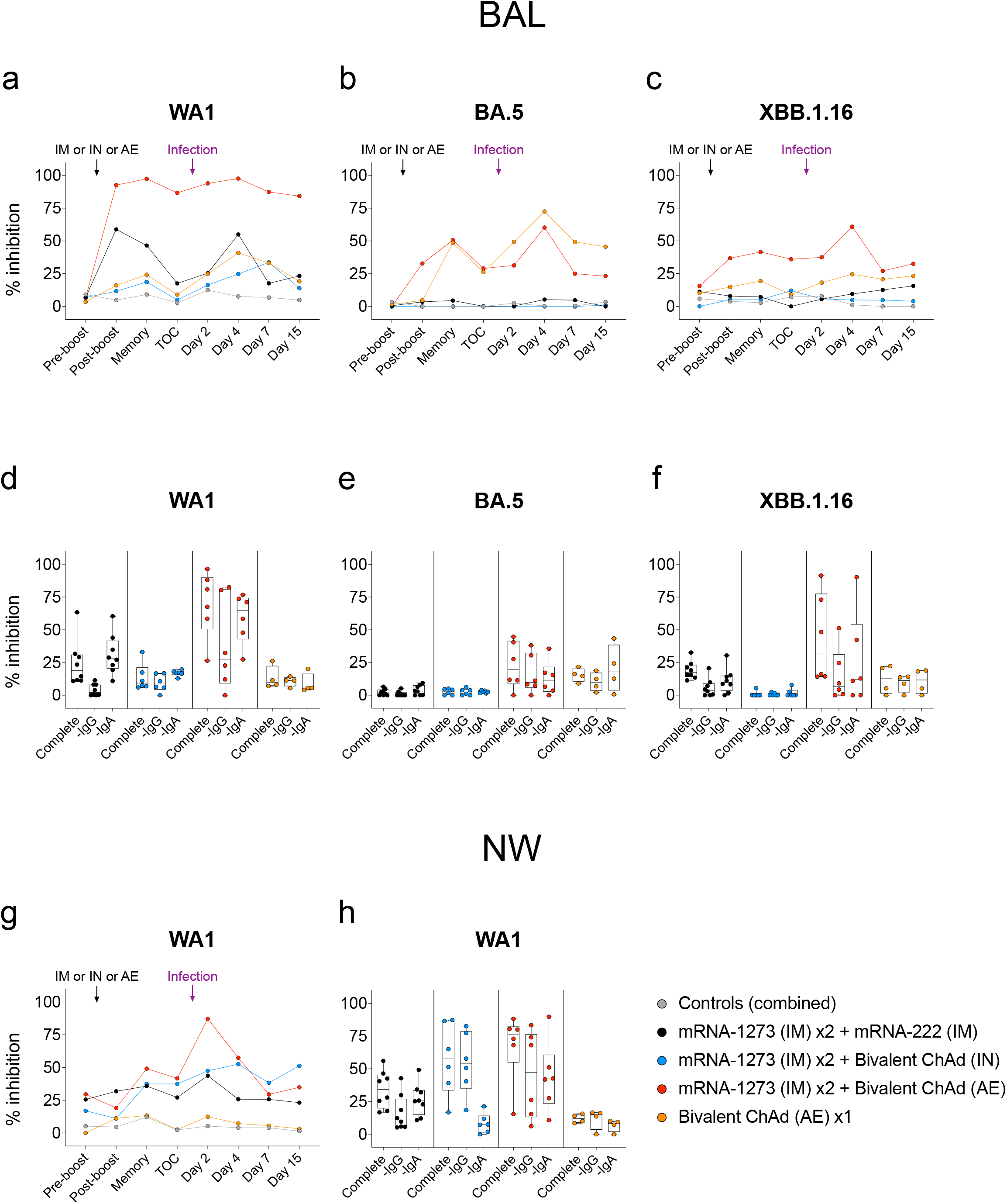
Functional IgG and IgA responses in the upper and lower airway following vaccination. NHP (*n*=4-8 per group) were administered mRNA-1273 or control mRNA at weeks 0 and 4 and boosted at week 32 with the indicated vaccine. (**a-c**) BAL and (**g**) NW were collected pre-boost (week 28), post-boost (week 34), at a memory timepoint (week 40), immediately prior to challenge (week 48) and post-challenge (days 2, 4, 7 and 15). WA1 (**a, g**), BA.5 (**b**) and XBB.1.16 (**c**) S binding to ACE2 was measured with and without the addition of mucosal fluids to determine percentage inhibition as a surrogate for functional antibodies. Symbols in **a-c** and **g** indicate median percent inhibition of each group. Mucosal fluids at a late memory timepoint pre-challenge (week 44) from BAL (**d-f**) or NW (**h**) were used to inhibit WA1 (**d, h**), BA.5 (**e**) or XBB.1.16 (**f**) S binding to ACE2 as complete fluid or after the selective depletion of either IgG or IgA. Symbols in **d-f** and **h** represent individual NHP. Boxes represent interquartile range with median indicated by solid line.

The AE boost increased ACE2 blocking antibodies in BAL fluid, with median binding inhibition against WA1 rising to 92.7% two weeks after the boost, and this activity remained stable over the subsequent 14 weeks. Inhibition of XBB.1.16 binding also increased (8% to 36.7%) (**Fig. 5c**). The AE prime group elicited ACE2 blocking antibody responses, although these were principally directed against BA.5 possibly reflecting the more immunogenic 6-P-stabilized structure of the BA.5 S compared to the 2-P-stabilized Wuhan S (**Fig. 5b**). In comparison, the IN boost did not appreciably increase ACE2 binding inhibition in BAL fluid against any variant.

Following XBB.1.16 challenge in the AE boosted group, there was an anamnestic response in the lungs with ACE2 binding inhibition for XBB.1.16 peaking at 60.9% on day 4 post-challenge (**Fig. 5c**), which was the same timepoint when we had observed a transient increase in BAL IgG binding titers (**Extended Data Fig. 4c**). We did not observe a corresponding increase in WA1-targeting functional antibodies, presumably because responses were already at the upper limit of the dynamic range prior to challenge. Anamnestic inhibitory antibody responses in the lung to WA1 were mounted by day 4 in the IM boosted and AE prime groups and by day 7 in the IN boosted group (**Fig. 5a**).

In the nose, pre-boost median ACE2 binding inhibition to WA1 was 21.3% and only marginally different two weeks after any boost (**Fig. 5g**). However, by the memory timepoint at week 40, binding inhibition had increased to 35.8%, 37.4% and 49.2% in the IM, IN and AE boost groups, respectively, reflecting different antibody induction kinetics in the upper and lower airways. Virus challenge elicited a recall response in all boosted groups by days 2-4, with peak median ACE2-binding inhibition values of 43.8%, 52.6% and 87.2% in the IM, IN and AE boost groups, respectively. It is notable that the IN and AE boosted groups, which had the greatest inhibitory responses in the NW at the memory timepoint and later at the time of challenge, were the only two groups to significantly suppress virus replication in the nose within two days of challenge.

Finally, we assessed the relative contributions of IgG and IgA to neutralization in mucosal fluids by measuring ACE2 binding inhibition after *in vitro* depletion of either IgG or IgA. Samples were collected at week 44, a post-boost memory timepoint. In BAL fluid, IgA depletion had limited effect on inhibitory responses in any group except for a modest reduction for XBB.1.16-targeting antibodies following an AE boost (**Fig. 5d-f**). However, IgG depletion reduced median WA1 S-ACE2 binding inhibition in BAL fluid from 18.9% to 2.2% in the IM boosted group and from 74.2% to 27.5% in the AE boosted group (**Fig. 5d**). All other groups showed limited binding inhibition.

In the upper airway, IgG depletion substantially reduced WA1 S-ACE2 binding inhibition in the IM boosted group from 34.3% to 10.9%, whereas IgA depletion had a lesser effect, reducing inhibition to 25.5% (**Fig. 5h**). For the AE boosted group, both IgG and IgA contributed to blockade of ACE2 binding, with inhibition declining from 76.5% to 47.1% and 42.2%, respectively. Importantly, the IN boosted group was unique in that inhibition was predominately IgA-mediated, as IgA depletion almost completely eliminated binding inhibition activity.

To summarize, the mRNA IM boost primarily elicited IgG responses in the blood, nose and BAL with limited IgA in mucosal sites. In contrast, the IN boost induced high titers of functional IgA in the upper airway and more modest titers of systemic IgG. The AE boost elicited IgG and IgA responses in blood, nose and BAL compartments.

### Mucosal ChAd-SARS-CoV-2-S vaccine expands antigen-specific T cells

IM administered mRNA or adenoviral vaccines induce CD4^+^ and CD8^+^ T cell responses against SARS-CoV-2, which may contribute to protection^27, 69, 70, 71, 72, 73^. Thus, we assessed the kinetics of T cell responses in blood and BAL after two-dose primary mRNA vaccination and boosting by the different IM and mucosal regimens or priming with adenoviral-vectored vaccine by an AE route. Following a boost with the IM mRNA vaccine or mucosal adenoviral-vectored vaccine, there were no further increases in the frequency of SARS-CoV-2-specific T_H_1, T_FH_ or CD8^+^ subsets in the blood above those elicited by the two-dose primary series (**Fig. 6a-d, Extended Data Fig 5**). However, in the BAL, there was an increase in S-specific T_H_1 and CD8^+^ T cells following the AE boost (**Fig. 6e, g**). The T cell response was highest in the AE prime group, which had the greatest frequency of S-specific T_H_1 and CD8^+^ T cells with median values of 14.1% and 4.7%, respectively at week 34 compared to 1.2% and 1.4% in the AE boost group, although the frequency of S-specific CD8^+^ T cells in the AE boost group did increase to 11.8% by week 48. For the IM, IN and control groups, the median frequencies of S-specific T_H_1 and CD8^+^ T cells in the BAL were all less than 1% each at both weeks 34 and 48. This suggests that antigen presentation in the lungs and MALT is an especially efficient way to induce T cell responses in the BAL.

**Figure 6.**
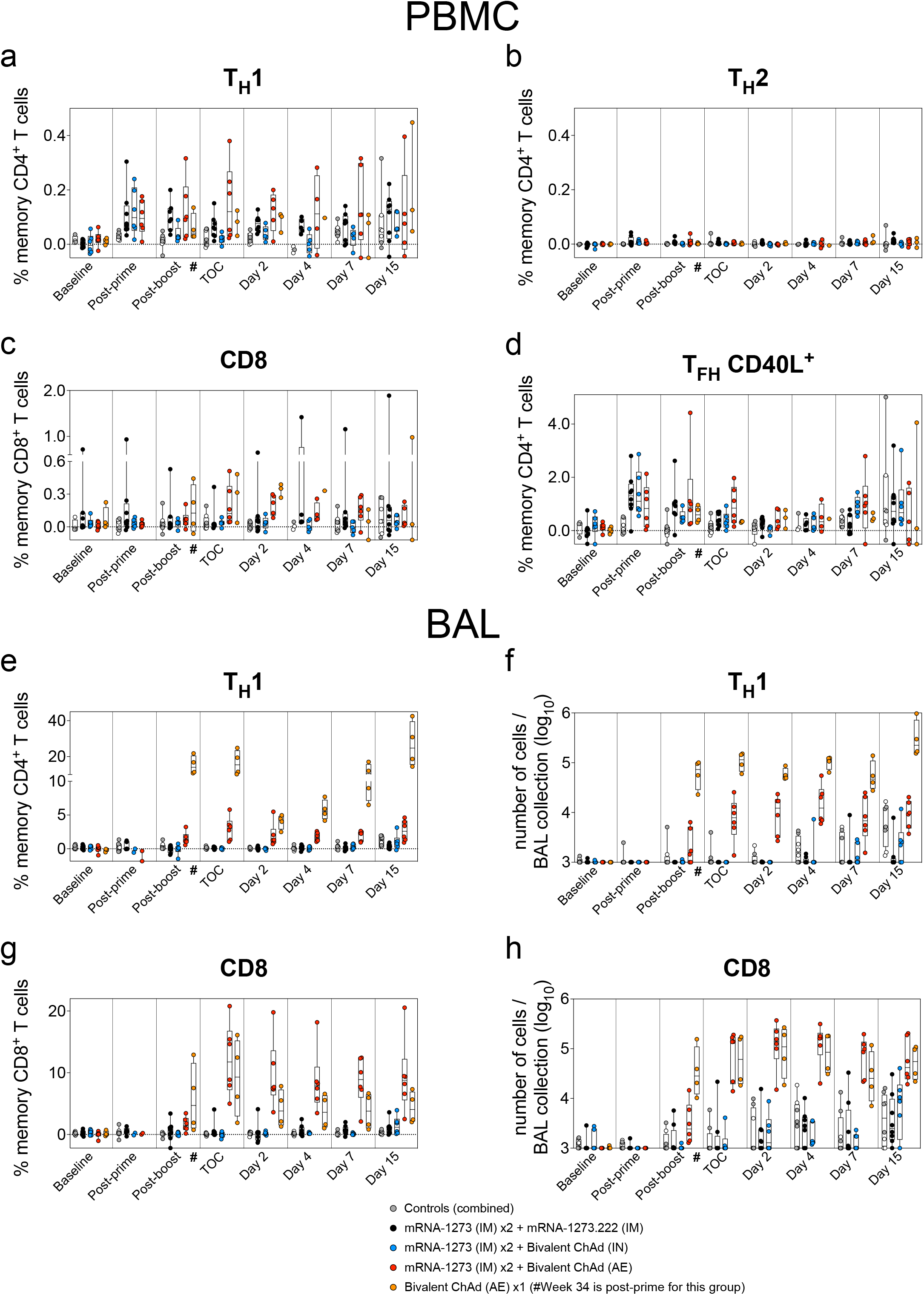
AE immunization elicits durable CD4 and CD8 T cell responses in the BAL. (**a-d**) PBMC and (**e-h**) BAL fluid were collected pre-vaccination (baseline) and at weeks 6 (post-prime), 34 (post-boost) and 48 (time of challenge) as well as on days 2, 4, 7 and 15 post-challenge. Lymphocytes were stimulated with SARS-CoV-2 S1 and S2 peptide pools (WA1) and then measured by intracellular staining. (**a-b, e**) Percentage of memory CD4^+^ T cells with (**a, e**) T_H_1 markers (IL-2, TNF-α or IFN-γ) or (**b**) T_H_2 markers (IL-4 or IL-13) following stimulation. (**c, g**) Percentage of memory CD8^+^ T cells expressing IL-2, TNF-α or IFN-γ following stimulation. (**d**) Percentage of T_FH_ cells that express CD40L following stimulation. Breaks in y-axis indicate a change in scale without a break in the range depicted. Dotted lines set at 0%. Absolute number of S-reactive (**f**) T_H_1 CD4^+^ or (**h**) CD8^+^ T cells in the BAL are also depicted. Counts below a value of 1 (due to background subtraction) were replaced with a value of 1 for data in **f** and **h**. Circles in **a-h** indicate individual NHP. Boxes represent interquartile range with the median denoted by a horizontal line. Reported values may be negative due to background subtraction and may extend below the range of the y-axis. # indicates that while sample collection for AE prime cohort (orange) occurred on week 34, week 34 was two weeks following the single AE prime rather than two weeks following a boost as in all other groups.

T cells may contribute to vaccine-mediated protection against severe SARS-CoV-2 disease in the lung^28, 54, 74^. Thus, we assessed the magnitude of the antigen-specific T cell compartment in the BAL following virus challenge. There was no significant increase in the number of S-specific T cells in the AE boost group at any time or in the AE prime group until day 15, at which time virus had already been cleared from the lungs; there was a trend towards a small increase in antigen-specific lung T cell counts in the IM and IN boost groups beginning at day 4 (**Fig 6f, h**). Responses to the N protein, which was not incorporated into the vaccine, were highest in the control group followed by the IM boost group (**Extended Data Fig 6**), whereas cohorts that received mucosal vaccines had low to undetectable N-specific T cell responses, consistent with rapid suppression of virus replication.

## Discussion

In this study, we evaluated whether targeting a SARS-CoV-2 vaccine directly to the mucosa could boost upper airway immune responses such that durable protection could be achieved against a heterologous highly transmissible immune-evading variant. To achieve optimal mucosal boosting, we compared two different delivery devices. Our principal findings were: (1) tissue-specific induction of airway mucosal antibodies is dependent on vaccine delivery route, with AE vaccines eliciting mucosal responses foremost in the lungs and secondarily in the nose, whereas IN vaccination elicited immunity primarily in the nose; (2) antigen-specific immunoglobulin class is dependent on site of antigen presentation, with IM vaccines primarily boosting IgG and mucosal vaccines inducing both IgG and IgA; (3) immunity deriving from mucosal adenoviral-vectored vaccines was durable over a period of 5 months in contrast to the rapid peak and subsequent waning typical of IM vaccination; and (4) control of virus in the lower airway, control of virus in the upper airway and control in both compartments were features of IM, IN and AE vaccination, respectively.

These data indicate that vaccines directed at both the lungs and nose (AE groups) can induce broad multi-compartment mucosal immunity, which effectively and rapidly suppresses virus replication in both the upper and lower respiratory tract such that insufficient antigen is available to promote systemic recall responses and would be consistent with preventing infection. However, mucosal vaccination directed primarily at the nose (IN boost group), while capable of boosting upper airway IgA titers and preventing local virus replication, did not suppress virus replication in the lungs as effectively as the AE boost. This highlights that the form of mucosal delivery can optimize immune responses in the specific upper and lower airway compartments and that this is associated with control of virus at those sites.

The prevention or substantive inhibition of most respiratory virus infections is mediated by antibodies in the upper and lower airways. For instance, against SARS-CoV-2, S-specific mucosal IgG antibodies are readily elicited by IM mRNA vaccination and correlate with protection^15, 46, 75^. By contrast, IM mRNA vaccines against SARS-CoV-2 elicit low to undetectable secretory IgA responses^26, 32, 66, 76^. Because IgA has been shown to correlate with protection from upper airway infection for SARS-CoV-2, influenza and respiratory syncytial virus^31, 32, 77, 78, 79, 80^, we determined whether mucosal vaccines would increase IgA production, and if this was associated with protection. Indeed, our findings show that IN delivery of adenoviral vaccines as a prime or boost induces upper airway IgA, which may be due to activation of IgA-producing plasmablasts and memory B cells in the nose and nasal associated lymphoid tissue (NALT)^81, 82, 83^. Mechanistically, the finding that IN boosting predominantly increased mucosal IgA levels, rather than IgG, suggests that local antigen production in the nose, whether from a vaccine or virus infection, primarily elicits IgA responses and that the IgG present in the upper airway derives from circulation by transudation^31, 32, 84^. Thus, enhanced IgA production by targeted vaccine delivery to the nasal mucosa from the IN boost could overcome the limited protection against current SARS-CoV-2 variants at that site conferred by IM-elicited neutralizing IgG^15, 16, 85^, which had previously been sufficient to confer protection against ancestral variants within a few months of immunization. It is notable that serum neutralizing antibody titers to circulating variants EG.5.1 and BA.2.86 were detectable following IM, but not IN, boosting and that these responses had no association with protection in the upper airway. By contrast, IN boosting essentially prevented virus replication in the upper airway with limited boosting of serum neutralizing titers; however, the association with higher IgA levels in the upper airways mediating functional neutralization suggest this as a potential mechanism of protection.

Virus-vectored vaccines not only elicit antibody immunity but are effective at inducing T cell responses^27, 35, 54, 69, 71^. In preclinical rodent and NHP models as well as correlative data from assessing responses from blood of humans, T cell immunity has been suggested to have a role in mediating protection against SARS-CoV-2, most notably against severe disease^28, 74, 86, 87^. We showed that AE-delivered ChAd-vectored vaccines induced high frequencies of CD4^+^ and CD8^+^ T cells in BAL that were sustained, which is consistent with prior studies in NHP using AE delivery of other antigens^88, 89, 90, 91^. Regarding the role of T cells in mediating protection in this study, it is notable that the frequency of S-specific T cells post-challenge did not significantly increase until after virus replication had been controlled in animals that were boosted by AE adenoviral vaccine or IM mRNA. Of note, reduction of virus replication in the BAL at day 2 post-challenge in animals boosted by IN vaccine was evident even though there were low to undetectable T cell responses. The modest T cell expansion in the BAL in any of the vaccine groups occurred with slower kinetics with the peak occurring near day 15 than for the anamnestic humoral responses, which peaked around day 4 consistent with prior studies showing early anamnestic antibody responses in BAL commensurate with virus control against SARS-CoV-2 in NHP^26, 67^. Together, these data further support a dominant antibody-mediated mechanism of protection after mucosal vaccine boosting both in the upper and lower airway. However, the contribution of T cells to protection cannot be fully elucidated without depletion studies and further investigation of such responses in the upper airway. Nevertheless, the induction of high frequency T cell responses in the lungs following AE boosting or priming could provide another layer of immunity in the context of further antigenic shift centered on B cell epitopes^92, 93, 94^.

A key aspect of optimizing mucosal immune responses in humans by vaccination will be the device used for delivery. We chose two different approaches selectively targeting the upper (IN) and lower airways (AE) using devices with extensive safety data in humans. The eFlow Technology used in the Investigational eFlow Nebulizer System (PARI) for AE delivery has been approved for treatment of cystic fibrosis^95^ and as shown here is highly efficient for generating robust antibody and T cell responses in the lung^88, 90, 91^. The MAD device used for IN delivery is similar to an IN sprayer used to administer the live attenuated quadrivalent influenza vaccine^96, 97^. Furthermore, the ChAdOx1 nCov-19 (AZD1222) vaccine, which uses a different adenovirus serotype than ChAd-SARS-CoV-2-S, was administered by the MAD device in a Phase I clinical trial for safety and immunogenicty^37^. In contrast to the data reported here, the ChAdOx1 nCov-19 clinical study showed limited mucosal IgA and IgG responses, limited peripheral T cell responses and little to no systemic humoral responses. These discordant findings may be due to one or more factors including: 1) the use of a different simian adenovirus strain with possibly distinct tropism; 2) a higher vaccine dose (1 × 10^11^ virus particles of ChAd-SARS-CoV-2-S as compared to 5 × 10^9^ - 5 × 10^10^ for ChAdOx1); and 3) the prefusion stabilization of S for ChAd-SARS-CoV-2-S as compared to wild-type S protein (used in ChAdOx1), as stabilization is more efficient at inducing neutralizing responses against class I fusion proteins^98, 99, 100^.

The selection of the optimal route of immunization and choice of a mucosal delivery device for limiting or preventing infection in humans may depend on how SARS-CoV-2 is transmitted. While virus is likely emitted as small aerosol droplets (<5 μm) that are generated deep in the lower airways^101, 102, 103, 104^, it remains unclear which anatomical site in the upper respiratory tract is the primary site of infection after natural exposure. While we did not achieve full protection in the lungs following IN boosting via MAD, it is notable that we experimentally administered our challenge stock via IN and IT routes. If natural infection occurs primarily in the nose, then additional delivery of challenge virus directly into the trachea in our study may have circumvented the ability of the IN vaccine to block the infection at its initial portal of entry.

### Limitations of the study

First, while we model the impact of prior immune exposure to multiple antigens (WA1, BA.5), we have not attempted to recapitulate the antecedent exposures that humans now have including infections before or after vaccination and additional vaccine doses. However, two mRNA vaccinations are likely sufficient to induce some level of immune imprinting. Second, while we included the AE prime group to characterize the role of prior immunity and the potential for use of a mucosal vaccine in a primary regimen, because of limited numbers of NHP available and due to the already large scope of the study, we could not include a similar IN prime arm. Likewise, we did not have a control group for IN adenovirus exposure using a vaccine that did not contain the SARS-CoV-2 antigen. As the challenge was done 18 weeks after boosting and the control AE group had no effect on protection, it is likely that this additional IN control would not have exerted any effect on virus replication. Finally, we did not administer ChAd-SARS-CoV-2-S via IM route, as the major comparison was with the currently available IM mRNA vaccines; however, a direct comparison of adenoviral-vectored vaccines delivered via IM and AE routes could allow us to better distinguish features of the enhanced immunogenicity that are due to route of administration.

In summary, this study provides a proof-of-principle for mucosal vaccination in a relevant pre-clinical model to achieve broad and durable cross-variant humoral and cellular immunity with functional prevention of XBB.1.16 infection. While we observed these results with an adenoviral-vectored vaccine, it is possible that alternative vaccine platforms, including live virus, protein or mRNA, could achieve similar results if delivered by a mucosal route. While there are rodent data for IN delivery of protein and mRNA^33, 34^, there are more limited NHP data modeling prior immunity by widely used mRNA vaccines prior to mucosal boosting to demonstrate durable protection against infection with a highly transmissible variant. Indeed, for NHP studies and likely for humans, protein and mRNA may need to be formulated appropriately to penetrate the upper airway mucosal tissue^105^, which is a natural feature of many vaccines derived from viruses that have evolved for this purpose. Furthermore, the choice of adenovirus serotype may be relevant as there may be distinct preferences for replication in the upper airway as compared to the gut^106^. The data presented here provide a roadmap for next-generation vaccines against COVID-19 and other respiratory pathogens with pandemic potential to achieve near-sterilizing prevention of infection, block transmission and potentially limit the rate of virus evolution. This approach also has the potential to reduce global virus burden and alter the outbreak dynamics of disease.

## Methods

### Rhesus macaque model

At the time of study enrollment, two- to six-year-old male Indian-origin rhesus macaques were primed with mRNA vaccine or were administered control mRNA. Animals were later stratified into groups for vaccine boosting based on age and weight. Animals were housed at Bioqual, Inc. (Rockville, MD) and all experiments were conducted according to NIH regulations and standards on the humane care and use of laboratory animals as well as the Animal Care and Use Committees of the NIH Vaccine Research Center and Bioqual, Inc.

### Preclinical mRNA and adenoviral-vectored vaccines

A sequence-optimized mRNA encoding prefusion-stabilized SARS-CoV-2 S protein containing 2 proline stabilization mutations (S-2P)^100, 107^ for Wuhan-1 or bivalent Wuhan-1/BA.5 was synthesized *in vitro* and formulated^108^. The ChAd-SARS-CoV-2-BA.5-S vector expressed a prefusion-stabilized S glycoprotein of BA.5 (GenBank: QJQ84760; T19I, L24S, del25-27, del69-70, G142D, V213G, G339D, S371F, S373P, S375F, T376A, D405N, R408S, K417N, N440K, G446S, L452R, S477N, T478K, E484A, F486V, Q498R, N501Y, Y505H, D614G, H655Y, N679K, P681H, N764K, D796Y, Q954H, N969K) containing six proline substitutions (F817P, A892P, A899P, A942P, K986P and V987P) and furin cleavage site substitutions (RRARS to GSASS, residues 682–686) as described elsewhere^109^. Control mRNA “NTFIX-01 (Not Translated Factor 9)” was synthesized and similarly formulated into lipid nanoparticles as previously described^48^.

The ChAd-SARS-CoV-2-S replication-incompetent vector (simian Ad36) encoding the prefusion-stabilized SARS-CoV-2 S-2P and empty ChAd-Control vector were generated as described previously^40^. The ChAd-SARS-CoV-2-BA.5-S genome was rescued following transfection of T-REx™-293 Cell Line (Invitrogen, R710-07). Replication-incompetent ChAd-SARS-CoV-2-BA.5-S, ChAd-SARS-CoV-2-S and ChAd-Control vectors were scaled up in HEK-293 cells (ATCC, CRL-1573) and purified by CsCl density-gradient ultracentrifugation. Viral particle concentrations were determined by spectrophotometry at 260 nm as described^110^.

### Vaccine delivery to rhesus macaques

IM delivery of mRNA vaccines or control mRNA was administered in 1 mL of formulated lipid nanoparticles diluted in phosphate-buffered saline (PBS) into the right quadricep as previously described^26, 46, 111^. For AE delivery, each animal was administered 1 mL ChAd-SARS-CoV-2-S or control ChAd (diluted in PBS to a concentration of 1×10^11^ vp) via pediatric silicone face mask (PARI SMARTMASK Baby/Kids) attached to an Investigational eFlow Nebulizer System (PARI) that delivered 4 μm particles deep into the lung of anesthetized macaques, as previously described^112^. For IN delivery, each animal was administered 200 µL of ChAd-SARS-CoV-2-S into each nostril for a total volume of 400 µL (diluted in PBS to a concentration of 2.5×10^11^ vp) via MAD Nasal™ (Teleflex). MAD was placed snugly against the nostril, aiming slightly up and outward toward the top of the ear. Mist emanating from MAD contained 30-100 μm particles deposited into the upper airway.

### Immunization and challenge schedule

20 NHP were primed at weeks 0 and 4 with two doses of 30 µg mRNA-1273 via IM route. At week 32 the following groups received a third dose: 1) 8 NHP (IM boost) received 30 µg of mRNA-1273.222 via IM route; 2) 6 NHP (IN boost) received 10^11^ vp of the bivalent cocktail of ChAd-SARS-CoV-2-Wuhan-1-S and ChAd-SARS-CoV-2-BA.5-S ^54^ via IN route; 3) 6 NHP (AE boost) received 10^11^ vp of the bivalent cocktail of ChAd-SARS-CoV-2-S via AE route. Also, at week 32, a naïve cohort of 4 NHP (AE prime) were administered a first dose of 10^11^ vp of the bivalent cocktail of ChAd-SARS-CoV-2-S via AE route.

Finally, an additional group of 8 NHP received two doses of 30 µg of control mRNA via IM route at weeks 0 and 4. At week 32, 4 of these NHP received 10^11^ vp of control ChAd via AE route.

All NHP were challenged at week 50 (18 weeks after final immunization) with a total dose of 3 × 10^5^ PFU of SARS-CoV-2 Omicron XBB.1.16. The virus inoculum was administered as 2.25 ×10^5^ PFU in 3mL via IT route and 0.75 ×10^5^ PFU in 1mL via IN route (MAD) with a volume of 0.5 mL distributed evenly into each nostril.

### Isolation and sequencing of XBB.1.16 challenge stock

XBB.1.16 (EPI_ISL_17417328) was isolated from a residual nasal swab kindly provided by Dr. Benjamin Pinsky (Stanford University). Virus was plaque purified and propagated once in VeroE6-TMPRSS2 cells to generate a working stock. XBB.1.16 stock was then sequenced as previously described^46, 47^. Briefly, NEBNext Ultra II RNA Prep reagents and multiplex oligos (New England Biolabs) were used to prepare Illumina-ready libraries, which were sequenced on a MiSeq (Illumina). Demultiplexed sequence reads were analyzed in the CLC Genomics Workbench v.23.0.1 by (1) trimming for quality, length, and adaptor sequence, (2) mapping to the Wuhan-Hu-1 SARS-CoV-2 reference (GenBank no. NC_045512), (3) improving the mapping by local realignment in areas containing insertions and deletions (indels) and (4) generating both a sample consensus sequence and a list of variants. Default settings were used for all tools.

### Cell lines

Cell lines used for ChAd production include T-REx™-293 Cell Line (Invitrogen, R710-07) for rescue of ChAd-SARS-CoV-2-BA.5-S genome and HEK-293 cells (ATCC, CRL-1573) for scaling up of ChAd vectors.

Cell lines used for propagation of XBB.1.16 challenge stock and TCID_50_ assays include VeroE6-TMPRSS2 and Vero-ACE2/TMPRSS2, respectively, which were provided by Adrian Creanga (VRC/NIAID). Both lines were cultured in complete DMEM medium consisting of 1x DMEM (VWR, #45000-304), 10% FBS, 2mM L-glutamine and 1x antibiotic as previously described^113^. 293T-human ACE2 cells (Drs. Michael Farzan and Huihui Mu at UF Scripps Institute) were used for pseudovirus neutralization assays. Additional experimental details contained in relevant method section below.

### Subgenomic RNA quantification

sgRNA was isolated and quantified by researchers blinded to vaccine status as previously described^46^. Briefly, total RNA was extracted from BAL fluid and NS using RNAzol BD column kit (Molecular Research Center). PCR reactions were conducted with TaqMan Fast Virus 1-Step Master Mix (Applied Biosystems), forward primer in the 5’ leader region and N gene-specific probe and reverse primer as previously described^47^:

sgLeadSARSCoV2_F: 5’-CGATCTCTTGTAGATCTGTTCTC-3’
N2_P: 5’-FAM-CGATCAAAACAACGTCGGCCCC-BHQ1-3’
wtN_R: 5’-GGTGAACCAAGACGCAGTAT-3’

Amplifications were performed with a QuantStudio 6 Pro Real-Time PCR System (Applied Biosystems). The assay lower LOD was 50 copies per reaction.

### TCID50 quantification

Infectivity of virus in NHP NS, NW and BAL was determined by TCID_50_ assay on Vero-ACE2/TMPRSS2. One hour prior to the assay, growth media was removed from the cells and replaced with 180 μL / well of 2% cDMEM (DMEM supplemented with 2% FBS, 2mM L-glutamine, and 1x antibiotic). Samples were diluted serially, 10-fold in DMEM containing 2% FBS. 20 μL diluted sample were added to cells in quadruplicate. After 5 days, infectious media was removed from the cells, which were then fixed and stained with crystal violet (20% methanol, 2.5 g crystal violet). Virus replication was scored as a lack of crystal violet staining. TCID_50_ values were calculated using the Reed-Muench method^114^. The lower LOD for quantification of virus titer was 108.

### Pseudovirus neutralization

The SARS-CoV-2 lentiviral pseudotyped neutralization assays were performed on integrated automation platforms consisting of a Biomek liquid handler from Beckman Coulter, ambient temperature labware hotel (Thermo Scientific), 37°C incubator (Thermo Scientific) and Molecular Devices Paradigm Multimode reader as previously described^47^. The automated assay methods were operated through Beckman Coulter SAMI EX software. On day 1, samples were diluted starting at 1:20 and then serially diluted 4-fold (7 times) in D10 culture medium (10% FBS, DMEM, 0.3 µL / mL puromycin) in a bulk sterile polypropylene 384-well deep-well plate. The diluted samples were transferred from bulk dilution plates into individual 384-well black tissue culture plates at 30 µL per well (Thermo Scientific Nunc 384-well polystyrene plates, cell culture surface; Catalog #164564). SARS-CoV-2 S-pseudotyped viruses were diluted in D10 and added at 30 µL per well into tissue culture plates containing the serially diluted samples, followed by 45 minutes incubation at 37°C w/ 5% CO_2_. 293T-human ACE2 reporter cells were added at a concentration of 10,000 cells per well in 20 µL into virus/sample tissue culture plates, followed by 72-hour incubation at 37°C w/ 5% CO_2_. On day 4 (72 hours from Day 1), 50 µL of culture medium was removed from the plates and 30 µL of luciferase substrate (Perkin Elmer Britelite Plus Catalog #6066769) was added. The plates were incubated at room temperature (RT) for 2 minutes and then samples/luciferase were mixed. The luminescence signal (RLU, relative luminescence unit) was measured using a Paradigm Multimode reader. The neutralization percentage (%) of test sample was determined by normalization of the test sample RLU to the RLU of virus and cell control wells with the following calculation: percentage (%) = [(test wells – average of cell control wells) - (average of virus control wells – average cell control wells)] ÷ (average virus control wells – average cell control wells) × 100. The neutralization curve fit was generated on a NAB analysis module on Labkey web-based server with 5 parameter non-linear regression. Neutralizing antibody titers are expressed as the reciprocal of the serum dilution required to reduce RLU by 50% and reported as inhibition dosage (ID_50_).

### Serum and mucosal antibody binding

Quantification of antibodies in the blood and mucosa were performed using multiplex electrochemiluminescence serology assays by Meso Scale Discovery Inc. (MSD) as previously described^47, 111^. Briefly, total IgG and IgA antigen-specific antibodies to variant SARS-CoV-2 S-were determined by MSD V-Plex SARS-CoV-2 Panel 36var3 for S (kindly provided by MSD) according to manufacturer’s instructions, except 25 μl of sample and detection antibody were used per well. Heat inactivated plasma was diluted 1:10000 in Diluent 100. BAL fluid and NW fluid were concentrated 10-fold with Amicon Ultra centrifugal filter devices (Millipore Sigma). Concentrated BAL samples were subsequently diluted 1:100 for IgG and 1:50 or 1:100 for IgA using Diluent 100 prior to measurement. Concentrated NW samples were subsequently diluted 1:10 or 1:25 for IgG and 1:50 or 1:100 for IgA using Diluent 100 prior to measurement. AU / mL were calculated for each sample using MSD reference standard curve with relevant reference for each antibody class and variant S except for anti-XBB.1.16 IgA titers, for which the SARS-CoV-2 WA1 reference standard was used.

### ACE2-S binding inhibition

ACE2 binding inhibition was performed as previously described^47^. Briefly, BAL and NW were concentrated 10-fold with Amicon Ultra centrifugal filter with 30kDa MWCO (Millipore). Concentrated samples were subsequently diluted 1:5 in Diluent 100 (MSD). The ACE2 binding inhibition assay for variant SARS-CoV-2 S (WA1 and BA.5) was performed with V-Plex SARS-CoV-2 Panel 32 (ACE2) Kit (MSD) per manufacturer’s instructions. Plates were read on MSD Sector S 600 instrument. Results are reported as % inhibition.

For XBB.1.16 variant, ACE2 binding inhibition was performed using a modified MSD platform assay. Briefly, after blocking MSD Streptavidin MULTI-ARRAY 384 well plates with Blocker A (MSD), the plates were coated with 1 μg / ml of biotinylated SARS-CoV-2 variant S-2P (XBB.1.16) and incubated for 1 hour at RT. The plates were then washed 5 times with wash buffer (1x PBS containing 0.05% Tween-20). Diluted samples were added to the coated plates and incubated for 1 hour at RT. MSD SULFO-TAG human ACE2 protein was diluted 1:200 and added to the plates. After 1 hour incubation at RT, the plates were washed 5 times with wash buffer and read on MSD Sector S 600 instrument after the addition of Gold Read Buffer B (MSD). Results are reported as % inhibition.

For depletion of specific antibody classes from concentrated NW and BAL (week 44), we used Pierce^TM^ Protein G magnetic beads (Thermo Fisher Scientific) according to manufacturer instructions. Briefly, 50 µL of NW or BAL was incubated for 2hrs at RT with 50 µL of Protein G magnetic beads previously equilibrated with PBS pH 7.4. After incubation, the beads were removed by magnetic stand and the flow-through was collected as IgG-depleted fluid. The magnetic beads were then washed 3x with PBS pH 7.4 and subsequently incubated with 100 µL of Pierce^TM^ IgG Elution Buffer, pH 2.0 (Thermo Fisher Scientific) for 10 min at RT. The beads were removed by magnetic stand and the eluant was collected and dialyzed against PBS pH 7.4 with Slide-A-Lyzer™ MINI Dialysis Devices, 20K MWCO (Thermo Fisher Scientific). The dialyzed eluant was collected as IgA-depleted fluid. Concentrations of IgG and IgA in the initial complete mucosal fluid, flow-through and eluant were determined by ELISA using the Isotyping Panel 1 NHP/Human Kit (MSD). An equal concentration of IgG and IgA for each sample was used in ACE2 binding inhibition assay as described above.

### Intracellular cytokine staining

Intracellular cytokine staining (ICS) was performed as previously described^26, 115^. Briefly, cryopreserved PBMC and BAL cells were thawed and rested overnight at 37°C / 5% CO_2_. After rest, cells were stimulated with SARS-CoV-2 S protein (S1 and S2) and nucleoprotein (N) peptide pools (JPT Peptides) at a final concentration of 2 μg / ml in the presence of 3 mM monensin for 6 hours. The S1, S2 and N peptide pools are comprised of 158, 157 and 102 individual peptides, respectively, as 15mers overlapping by 11 amino acids in 100% DMSO. Negative controls received an equal concentration of DMSO to that of peptide pools (final concentration of 0.5%). The following monoclonal antibodies were used: CD3 APC-Cy7 (clone SP34.2, BD Biosciences), CD4 PE-Cy5.5 (clone SK3, Thermo Fisher), CD8 BV570 (clone RPA-T8, BioLegend), CD45RA PE-Cy5 (clone 5H9, BD Biosciences), CCR7 BV650 (clone G043H7, BioLegend), CXCR5 PE (clone MU5UBEE, Thermo Fisher), CXCR3 BV711 (clone 1C6/CXCR3, BD Biosciences), PD-1 BUV737 (clone EH12.1, BD Biosciences), ICOS Pe-Cy7 (clone C398.4A, BioLegend), CD69 ECD (cloneTP1.55.3, Beckman Coulter), IFN-γ Ax700 (clone B27, BioLegend), IL-2 BV750 (clone MQ1-17H12, BD Biosciences), IL-4 BB700 (clone MP4-25D2, BD Biosciences), TNF-FITC (clone Mab11, BD Biosciences), IL-13 BV421 (clone JES10-5A2, BD Biosciences), IL-17 BV605 (clone BL168, BioLegend), IL-21 Ax647 (clone 3A3-N2.1, BD Biosciences) and CD154 BV785 (clone 24-31, BioLegend). Aqua live/dead fixable dead cell stain kit (Thermo Fisher Scientific) was used to exclude dead cells. All antibodies were previously titrated to determine the optimal concentration. Samples were acquired on a BD FACSymphony flow cytometer and analyzed using FlowJo version 10.9.0 (Treestar, Inc., Ashland, OR).

### Statistical analysis

For statistical analyses of virus titers (sgRNA and TCID_50_) in Figure 2, the three groups given adenoviral-vectored vaccine were compared against the pooled control arms at each timepoint. For each comparison, Kruskal-Wallis tests were used to compare all groups simultaneously at α=0.05, followed by comparisons for each vaccinated group and the pooled controls using Wilcoxon Rank-Sum tests at α=0.05/2 if the overall test was statistically significant. Reported pairwise *p* values were doubled to account for this adjustment. For analysis, values of sgRNA below the LOD were set at the LOD (50 copies) and culturable virus titers that were not determined or undetectable were set to the assay LOD of 108. *p* values are indicated by asterisks in the figures, and the sample *n* is listed in corresponding figure legends.

Humoral, cellular and virus assays were log-transformed as appropriate and reported as geometric means with error bars depicting geometric standard deviation as indicated. There are no adjustments for multiple comparisons, so all *p* values should be interpreted as suggestive rather than conclusive. All analyses are conducted using R version 4.1.0 unless otherwise specified.

## Data availability

All data are available in the main text or in the Extended Data figures.

## Acknowledgements

We would like to acknowledge Gabriela Alvarado, who provided organizational support. Alan Hoofring designed the graphical abstract. Manjula Basappa helped culture pseudovirus stocks for neutralization assays. We also are grateful for the work of the Nonhuman Primate Immunogenicity Core of the Vaccine Research Center (VRC) including members Briana Briscoe-Whatley, Mitzi Donaldson, Josue Marquez, Samantha Provost and Heather Tapley for processing the blood and tissue samples for this study. John Graves III, Ruth Woodward and Cole Honeycutt provided significant advice or support with regards to vaccination and sampling, and in the case of John Graves III, direct assistance. We would like to thank PARI for the Investigational eFlow Nebulizer System and silicone face mask. Finally, Sayda Elbashir and Darin Edwards (Moderna) kindly provided mRNA constructs and informed study design.

Funding was primarily provided by the Intramural Research Program of the VRC of the National Institute of Allergy and Infectious Diseases (NIAID) of the National Institutes of Health (NIH). Funding was also provided under HHSN272201400004C (NIAID Centers of Excellence for Influenza Research and Surveillance, CEIRS) and NIH P51 OD011132 awarded to Emory University. This work was also supported in part by the Emory Executive Vice President for Health Affairs Synergy Fund award, COVID-Catalyst-I3 Funds from the Woodruff Health Sciences Center and Emory School of Medicine, the Pediatric Research Alliance Center for Childhood Infections and Vaccines and Children’s Healthcare of Atlanta, and Woodruff Health Sciences Center 2020 COVID-19 CURE Award. Research support was provided to M.S.D. under NIH R01 AI157155 and the NIAID Centers of Excellence for Influenza Research and Response (CEIRR) contracts HHSN272201400008C, 75N93021C00014 and 75N93019C00051. Support to D.T.C. was provided by NIH R01 CA211096.

## Author information

### Contributions

M.G., J.M.T., M.R., M.S.D., D.C.D and R.A.S. designed experiments. M.G., B.J.F., S.F.A., D.R.F., A.M., E.L., M.E.D-G., M.R.B., L.A.S., B.C.L., Z.E.Z., E.M., R.C., M.N., Y.J., N.A.D-R., S.O., J.M., B.W., M.E., A.V.R., M.G.L., H.A., K.E.F., M.S.S., D.C.D and R.A.S. performed, analyzed and / or supervised in vitro experiments. L.P., J.M.T., A.D., K.K., S.Z., M.G.L. and H.A. performed and / or supervised NHP handling, sampling and immunizations. J.B.C. designed adenoviral constructs. I.P.D., E.A.K. and B.Y. cloned, produced and / or tested adenoviral constructs. D.T.C. and M.S.D. supervised adenoviral construct production. I.T., L.W., Q.Z. and P.D.K. provided and / or supervised production of other critical reagents. S.G., F.L. and A.R.H. sequenced and / or analyzed challenge stock genome. M.C.N. performed statistical analyses. Funding was provided by D.T.C., M.S.S., M.S.D. and the Intramural Research Program of the VRC. M.G., D.C.D and R.A.S. designed figures. M.G., M.S.D., D.C.D. and R.A.S. wrote manuscript. All authors reviewed manuscript and provided experimental feedback.

### Corresponding authors

Correspondence to Daniel C. Douek and Robert A. Seder.

## References

1. Andrews, N. et al. Duration of Protection against Mild and Severe Disease by Covid-19 Vaccines. N Engl J Med 386, 340–350 (2022).

2. Tseng, H.F. et al. Effectiveness of mRNA-1273 vaccination against SARS-CoV-2 omicron subvariants BA.1, BA.2, BA.2.12.1, BA.4, and BA.5. Nat Commun 14, 189 (2023).

3. Buchan, S.A. et al. Estimated Effectiveness of COVID-19 Vaccines Against Omicron or Delta Symptomatic Infection and Severe Outcomes. JAMA Netw Open 5, e2232760 (2022).

4. Tartof, S.Y. et al. Effectiveness of mRNA BNT162b2 COVID-19 vaccine up to 6 months in a large integrated health system in the USA: a retrospective cohort study. Lancet 398, 1407–1416 (2021).

5. Lin, D.Y. et al. Durability of Bivalent Boosters against Omicron Subvariants. N Engl J Med 388, 1818–1820 (2023).

6. Bowe, B., Xie, Y. & Al-Aly, Z. Acute and postacute sequelae associated with SARS-CoV-2 reinfection. Nat Med 28, 2398–2405 (2022).

7. Bahremand, T. et al. COVID-19 hospitalisations in immunocompromised individuals in the Omicron era: a population-based observational study using surveillance data in British Columbia, Canada. Lancet Reg Health Am 20, 100461 (2023).

8. Malahe, S.R.K. et al. Clinical Characteristics and Outcomes of Immunocompromised Patients With Coronavirus Disease 2019 Caused by the Omicron Variant: A Prospective, Observational Study. Clin Infect Dis 76, e172–e178 (2023).

9. Griggs, E.P. et al. Clinical epidemiology and risk factors for critical outcomes among vaccinated and unvaccinated adults hospitalized with COVID-19-VISION Network, 10 States, June 2021-March 2023. Clin Infect Dis (2023).

10. Starkey, T. et al. A population-scale temporal case-control evaluation of COVID-19 disease phenotype and related outcome rates in patients with cancer in England (UKCCP). Sci Rep 13, 11327 (2023).

11. Goldberg, Y. et al. Waning Immunity after the BNT162b2 Vaccine in Israel. N Engl J Med 385, e85 (2021).

12. Baden, L.R. et al. Phase 3 Trial of mRNA-1273 during the Delta-Variant Surge. N Engl J Med 385, 2485–2487 (2021).

13. Bergwerk, M. et al. Covid-19 Breakthrough Infections in Vaccinated Health Care Workers. N Engl J Med 385, 1474–1484 (2021).

14. Menegale, F. et al. Evaluation of Waning of SARS-CoV-2 Vaccine-Induced Immunity: A Systematic Review and Meta-analysis. JAMA Netw Open 6, e2310650 (2023).

15. Corbett, K.S. et al. Immune correlates of protection by mRNA-1273 vaccine against SARS-CoV-2 in nonhuman primates. Science, eabj0299 (2021).

16. He, X. et al. Low-dose Ad26.COV2.S protection against SARS-CoV-2 challenge in rhesus macaques. Cell 184, 3467–3473 e3411 (2021).

17. Rossler, A. et al. Characterizing SARS-CoV-2 neutralization profiles after bivalent boosting using antigenic cartography. Nat Commun 14, 5224 (2023).

18. Garcia-Beltran, W.F. et al. Multiple SARS-CoV-2 variants escape neutralization by vaccine-induced humoral immunity. Cell 184, 2372–2383 e2379 (2021).

19. Cromer, D. et al. Neutralising antibody titres as predictors of protection against SARS-CoV-2 variants and the impact of boosting: a meta-analysis. Lancet Microbe 3, e52–e61 (2022).

20. Carabelli, A.M. et al. SARS-CoV-2 variant biology: immune escape, transmission and fitness. Nat Rev Microbiol 21, 162–177 (2023).

21. Hacisuleyman, E. et al. Vaccine Breakthrough Infections with SARS-CoV-2 Variants. N Engl J Med 384, 2212–2218 (2021).

22. Reynolds, C.J. et al. Immune boosting by B.1.1.529 (Omicron) depends on previous SARS-CoV-2 exposure. Science 377, eabq1841 (2022).

23. Roltgen, K. et al. Immune imprinting, breadth of variant recognition, and germinal center response in human SARS-CoV-2 infection and vaccination. Cell 185, 1025–1040 e1014 (2022).

24. Alsoussi, W.B. et al. SARS-CoV-2 Omicron boosting induces de novo B cell response in humans. Nature 617, 592–598 (2023).

25. Mackin, S.R. et al. Fc-gammaR-dependent antibody effector functions are required for vaccine-mediated protection against antigen-shifted variants of SARS-CoV-2. Nat Microbiol 8, 569–580 (2023).

26. Gagne, M. et al. Protection from SARS-CoV-2 Delta one year after mRNA-1273 vaccination in rhesus macaques coincides with anamnestic antibody response in the lung. Cell 185, 113–130 e115 (2022).

27. Liu, J., et al. CD8 T cells contribute to vaccine protection against SARS-CoV-2 in macaques. Sci Immunol 7, eabq7647 (2022).

28. Rydyznski Moderbacher, C., et al. Antigen-Specific Adaptive Immunity to SARS-CoV-2 in Acute COVID-19 and Associations with Age and Disease Severity. Cell 183, 996–1012 e1019 (2020).

29. Knisely, J.M. et al. Mucosal vaccines for SARS-CoV-2: scientific gaps and opportunities-workshop report. NPJ Vaccines 8, 53 (2023).

30. Mostaghimi, D., Valdez, C.N., Larson, H.T., Kalinich, C.C. & Iwasaki, A. Prevention of host-to-host transmission by SARS-CoV-2 vaccines. Lancet Infect Dis 22, e52–e58 (2022).

31. Havervall, S. et al. Anti-Spike Mucosal IgA Protection against SARS-CoV-2 Omicron Infection. N Engl J Med 387, 1333–1336 (2022).

32. Zuo, F., Marcotte, H., Hammarstrom, L. & Pan-Hammarstrom, Q. Mucosal IgA against SARS-CoV-2 Omicron Infection. N Engl J Med 387, e55 (2022).

33. Mao, T. et al. Unadjuvanted intranasal spike vaccine elicits protective mucosal immunity against sarbecoviruses. Science 378, eabo2523 (2022).

34. Baldeon Vaca, G., et al. Intranasal mRNA-LNP vaccination protects hamsters from SARS-CoV-2 infection. Sci Adv 9, eadh1655 (2023).

35. Le Nouen, C. et al. Intranasal pediatric parainfluenza virus-vectored SARS-CoV-2 vaccine is protective in monkeys. Cell 185, 4811–4825 e4817 (2022).

36. Ponce-de-Leon, S. et al. Interim safety and immunogenicity results from an NDV-based COVID-19 vaccine phase I trial in Mexico. NPJ Vaccines 8, 67 (2023).

37. Madhavan, M. et al. Tolerability and immunogenicity of an intranasally-administered adenovirus-vectored COVID-19 vaccine: An open-label partially-randomised ascending dose phase I trial. EBioMedicine 85, 104298 (2022).

38. Wu, S. et al. Safety, tolerability, and immunogenicity of an aerosolised adenovirus type-5 vector-based COVID-19 vaccine (Ad5-nCoV) in adults: preliminary report of an open-label and randomised phase 1 clinical trial. Lancet Infect Dis 21, 1654–1664 (2021).

39. Singh, C. et al. Phase III Pivotal comparative clinical trial of intranasal (iNCOVACC) and intramuscular COVID 19 vaccine (Covaxin((R))). NPJ Vaccines 8, 125 (2023).

40. Hassan, A.O. et al. A Single-Dose Intranasal ChAd Vaccine Protects Upper and Lower Respiratory Tracts against SARS-CoV-2. Cell 183, 169–184 e113 (2020).

41. Hassan, A.O. et al. A single intranasal dose of chimpanzee adenovirus-vectored vaccine protects against SARS-CoV-2 infection in rhesus macaques. Cell Rep Med 2, 100230 (2021).

42. Bricker, T.L. et al. A single intranasal or intramuscular immunization with chimpanzee adenovirus-vectored SARS-CoV-2 vaccine protects against pneumonia in hamsters. Cell Rep 36, 109400 (2021).

43. Ying, B. et al. A bivalent ChAd nasal vaccine protects against SARS-CoV-2 BQ.1.1 and XBB.1.5 infection and disease in mice and hamsters. bioRxiv, 2023.2005.2004.539332 (2023).

44. Hassan, A.O. et al. An intranasal vaccine durably protects against SARS-CoV-2 variants in mice. Cell Rep 36, 109452 (2021).

45. Munster, V.J. et al. Respiratory disease in rhesus macaques inoculated with SARS-CoV-2. Nature 585, 268–272 (2020).

46. Corbett, K.S. et al. mRNA-1273 protects against SARS-CoV-2 beta infection in nonhuman primates. Nat Immunol 22, 1306–1315 (2021).

47. Gagne, M. et al. mRNA-1273 or mRNA-Omicron boost in vaccinated macaques elicits similar B cell expansion, neutralizing responses, and protection from Omicron. Cell 185, 1556–1571 e1518 (2022).

48. Corbett, K.S. et al. Protection against SARS-CoV-2 Beta variant in mRNA-1273 vaccine-boosted nonhuman primates. Science 374, 1343–1353 (2021).

49. Chandrashekar, A. et al. Vaccine Protection Against the SARS-CoV-2 Omicron Variant in Macaques. bioRxiv, 2022.2002.2006.479285 (2022).

50. Yu, J. et al. Protective efficacy of Ad26.COV2.S against SARS-CoV-2 B.1.351 in macaques. Nature 596, 423–427 (2021).

51. Routhu, N.K., et al. Efficacy of mRNA-1273 and Novavax ancestral or BA.1 spike booster vaccines against SARS-CoV-2 BA.5 infection in non-human primates. Sci Immunol, eadg7015 (2023).

52. Solforosi, L. et al. Booster with Ad26.COV2.S or Omicron-adapted vaccine enhanced immunity and efficacy against SARS-CoV-2 Omicron in macaques. Nat Commun 14, 1944 (2023).

53. Schiepers, A. et al. Molecular fate-mapping of serum antibody responses to repeat immunization. Nature 615, 482–489 (2023).

54. Ying, B., et al. A bivalent ChAd nasal vaccine protects against SARS-CoV-2 BQ.1.1 and XBB.1.5 infection and disease in mice and hamsters. bioRxiv (2023).

55. Li, D. et al. Breadth of SARS-CoV-2 neutralization and protection induced by a nanoparticle vaccine. Nat Commun 13, 6309 (2022).

56. Tseng, H.F. et al. Effectiveness of mRNA-1273 against SARS-CoV-2 Omicron and Delta variants. Nat Med 28, 1063–1071 (2022).

57. Bar-On, Y.M. et al. Protection by a Fourth Dose of BNT162b2 against Omicron in Israel. N Engl J Med 386, 1712–1720 (2022).

58. Suryadevara, N. et al. Neutralizing and protective human monoclonal antibodies recognizing the N-terminal domain of the SARS-CoV-2 spike protein. Cell 184, 2316–2331 e2315 (2021).

59. Winkler, E.S. et al. Human neutralizing antibodies against SARS-CoV-2 require intact Fc effector functions for optimal therapeutic protection. Cell 184, 1804–1820 e1816 (2021).

60. Schafer, A. et al. Antibody potency, effector function, and combinations in protection and therapy for SARS-CoV-2 infection in vivo. J Exp Med 218 (2021).

61. Levin, E.G. et al. Waning Immune Humoral Response to BNT162b2 Covid-19 Vaccine over 6 Months. N Engl J Med 385, e84 (2021).

62. Pegu, A. et al. Durability of mRNA-1273 vaccine-induced antibodies against SARS-CoV-2 variants. Science (2021).

63. Barouch, D.H. et al. Durable Humoral and Cellular Immune Responses 8 Months after Ad26.COV2.S Vaccination. N Engl J Med 385, 951–953 (2021).

64. Collier, A.Y. et al. Differential Kinetics of Immune Responses Elicited by Covid-19 Vaccines. N Engl J Med 385, 2010–2012 (2021).

65. Bladh, O. et al. Mucosal immune responses following a fourth SARS-CoV-2 vaccine dose. Lancet Microbe 4, e488 (2023).

66. Liew, F. et al. SARS-CoV-2-specific nasal IgA wanes 9 months after hospitalisation with COVID-19 and is not induced by subsequent vaccination. EBioMedicine 87, 104402 (2023).

67. Francica, J.R. et al. Protective antibodies elicited by SARS-CoV-2 spike protein vaccination are boosted in the lung after challenge in nonhuman primates. Sci Transl Med 13 (2021).

68. Piccoli, L. et al. Mapping Neutralizing and Immunodominant Sites on the SARS-CoV-2 Spike Receptor-Binding Domain by Structure-Guided High-Resolution Serology. Cell 183, 1024–1042 e1021 (2020).

69. Mercado, N.B. et al. Single-shot Ad26 vaccine protects against SARS-CoV-2 in rhesus macaques. Nature 586, 583–588 (2020).

70. Mateus, J. et al. Low-dose mRNA-1273 COVID-19 vaccine generates durable memory enhanced by cross-reactive T cells. Science 374, eabj9853 (2021).

71. Swanson, P.A., 2nd et al. AZD1222/ChAdOx1 nCoV-19 vaccination induces a polyfunctional spike protein-specific T(H)1 response with a diverse TCR repertoire. Sci Transl Med 13, eabj7211 (2021).

72. Tarke, A. et al. SARS-CoV-2 vaccination induces immunological T cell memory able to cross-recognize variants from Alpha to Omicron. Cell 185, 847–859 e811 (2022).

73. Egri, N. et al. Cellular and humoral responses after second and third SARS-CoV-2 vaccinations in patients with autoimmune diseases treated with rituximab: specific T cell immunity remains longer and plays a protective role against SARS-CoV-2 reinfections. Front Immunol 14, 1146841 (2023).

74. Wherry, E.J. & Barouch, D.H. T cell immunity to COVID-19 vaccines. Science 377, 821–822 (2022).

75. Tang, J., et al. Respiratory mucosal immunity against SARS-CoV-2 after mRNA vaccination. Sci Immunol 7, eadd4853 (2022).

76. Sheikh-Mohamed, S. et al. Systemic and mucosal IgA responses are variably induced in response to SARS-CoV-2 mRNA vaccination and are associated with protection against subsequent infection. Mucosal Immunol 15, 799–808 (2022).

77. Zohar, T. et al. Upper and lower respiratory tract correlates of protection against respiratory syncytial virus following vaccination of nonhuman primates. Cell Host Microbe 30, 41–52 e45 (2022).

78. Seibert, C.W. et al. Recombinant IgA is sufficient to prevent influenza virus transmission in guinea pigs. J Virol 87, 7793–7804 (2013).

79. Ambrose, C.S., Wu, X., Jones, T. & Mallory, R.M. The role of nasal IgA in children vaccinated with live attenuated influenza vaccine. Vaccine 30, 6794–6801 (2012).

80. Gould, V.M.W. et al. Nasal IgA Provides Protection against Human Influenza Challenge in Volunteers with Low Serum Influenza Antibody Titre. Front Microbiol 8, 900 (2017).

81. Shimoda, M. et al. Isotype-specific selection of high affinity memory B cells in nasal-associated lymphoid tissue. J Exp Med 194, 1597–1607 (2001).

82. Sanchez Montalvo, A., Gohy, S., Rombaux, P., Pilette, C. & Hox, V. The Role of IgA in Chronic Upper Airway Disease: Friend or Foe? Front Allergy 3, 852546 (2022).

83. Hodge, L.M. et al. Immunoglobulin A (IgA) responses and IgE-associated inflammation along the respiratory tract after mucosal but not systemic immunization. Infect Immun 69, 2328–2338 (2001).

84. Wagner, D.K. et al. Analysis of immunoglobulin G antibody responses after administration of live and inactivated influenza A vaccine indicates that nasal wash immunoglobulin G is a transudate from serum. J Clin Microbiol 25, 559–562 (1987).

85. Wellford, S.A. et al. Mucosal plasma cells are required to protect the upper airway and brain from infection. Immunity 55, 2118–2134 e2116 (2022).

86. Swadling, L. et al. Pre-existing polymerase-specific T cells expand in abortive seronegative SARS-CoV-2. Nature 601, 110–117 (2022).

87. Bertoletti, A., Le Bert, N. & Tan, A.T. SARS-CoV-2-specific T cells in the changing landscape of the COVID-19 pandemic. Immunity 55, 1764–1778 (2022).

88. Darrah, P.A. et al. Boosting BCG with proteins or rAd5 does not enhance protection against tuberculosis in rhesus macaques. NPJ Vaccines 4, 21 (2019).

89. White, A.D. et al. Spore-FP1 tuberculosis mucosal vaccine candidate is highly protective in guinea pigs but fails to improve on BCG-conferred protection in non-human primates. Front Immunol 14, 1246826 (2023).

90. Darrah, P.A. et al. Aerosol vaccination with AERAS-402 elicits robust cellular immune responses in the lungs of rhesus macaques but fails to protect against high-dose Mycobacterium tuberculosis challenge. J Immunol 193, 1799–1811 (2014).

91. Hokey, D.A. et al. A nonhuman primate toxicology and immunogenicity study evaluating aerosol delivery of AERAS-402/Ad35 vaccine: Evidence for transient t cell responses in peripheral blood and robust sustained responses in the lungs. Hum Vaccin Immunother 10, 2199–2210 (2014).

92. Tarke, A. et al. Impact of SARS-CoV-2 variants on the total CD4(+) and CD8(+) T cell reactivity in infected or vaccinated individuals. Cell Rep Med 2, 100355 (2021).

93. Riou, C. et al. Escape from recognition of SARS-CoV-2 variant spike epitopes but overall preservation of T cell immunity. Sci Transl Med 14, eabj6824 (2022).

94. Choi, S.J. et al. T cell epitopes in SARS-CoV-2 proteins are substantially conserved in the Omicron variant. Cell Mol Immunol 19, 447–448 (2022).

95. Sawicki, G.S., Chou, W., Raimundo, K., Trzaskoma, B. & Konstan, M.W. Randomized trial of efficacy and safety of dornase alfa delivered by eRapid nebulizer in cystic fibrosis patients. J Cyst Fibros 14, 777–783 (2015).

96. Block, S.L., Yi, T., Sheldon, E., Dubovsky, F. & Falloon, J. A randomized, double-blind noninferiority study of quadrivalent live attenuated influenza vaccine in adults. Vaccine 29, 9391–9397 (2011).

97. Block, S.L. et al. Immunogenicity and safety of a quadrivalent live attenuated influenza vaccine in children. Pediatr Infect Dis J 31, 745–751 (2012).

98. Chang, L.A. et al. A prefusion-stabilized RSV F subunit vaccine elicits B cell responses with greater breadth and potency than a postfusion F vaccine. Sci Transl Med 14, eade0424 (2022).

99. McLellan, J.S. et al. Structure-based design of a fusion glycoprotein vaccine for respiratory syncytial virus. Science 342, 592–598 (2013).

100. Pallesen, J. et al. Immunogenicity and structures of a rationally designed prefusion MERS-CoV spike antigen. Proc Natl Acad Sci U S A 114, E7348–E7357 (2017).

101. Alsved, M. et al. Size distribution of exhaled aerosol particles containing SARS-CoV-2 RNA. Infect Dis (Lond) 55, 158–163 (2023).

102. Groma, V. et al. Size distribution and relationship of airborne SARS-CoV-2 RNA to indoor aerosol in hospital ward environments. Sci Rep 13, 3566 (2023).

103. Hawks, S.A. et al. Infectious SARS-CoV-2 Is Emitted in Aerosol Particles. mBio 12, e0252721 (2021).

104. Wang, C.C. et al. Airborne transmission of respiratory viruses. Science 373 (2021).

105. Hartwell, B.L. et al. Intranasal vaccination with lipid-conjugated immunogens promotes antigen transmucosal uptake to drive mucosal and systemic immunity. Sci Transl Med 14, eabn1413 (2022).

106. Custers, J. et al. Vaccines based on replication incompetent Ad26 viral vectors: Standardized template with key considerations for a risk/benefit assessment. Vaccine 39, 3081–3101 (2021).

107. Wrapp, D. et al. Cryo-EM structure of the 2019-nCoV spike in the prefusion conformation. Science 367, 1260–1263 (2020).

108. Hassett, K.J. et al. Optimization of Lipid Nanoparticles for Intramuscular Administration of mRNA Vaccines. Mol Ther Nucleic Acids 15, 1–11 (2019).

109. Hsieh, C.L. et al. Structure-based design of prefusion-stabilized SARS-CoV-2 spikes. Science 369, 1501–1505 (2020).

110. Maizel, J.V., Jr., White, D.O. & Scharff, M.D. The polypeptides of adenovirus. I. Evidence for multiple protein components in the virion and a comparison of types 2, 7A, and 12. Virology 36, 115–125 (1968).

111. Corbett, K.S. et al. Evaluation of the mRNA-1273 Vaccine against SARS-CoV-2 in Nonhuman Primates. N Engl J Med 383, 1544–1555 (2020).

112. Song, K. et al. Genetic immunization in the lung induces potent local and systemic immune responses. Proc Natl Acad Sci U S A 107, 22213–22218 (2010).

113. Edara, V.V. et al. Infection- and vaccine-induced antibody binding and neutralization of the B.1.351 SARS-CoV-2 variant. Cell Host Microbe 29, 516–521.e513 (2021).

114. Lei, C., Yang, J., Hu, J. & Sun, X. On the Calculation of TCID50 for Quantitation of Virus Infectivity. Virologica Sinica 36, 141–144 (2021).

115. Donaldson, M.M., Kao, S.F. & Foulds, K.E. OMIP-052: An 18-Color Panel for Measuring Th1, Th2, Th17, and Tfh Responses in Rhesus Macaques. Cytometry A 95, 261–263 (2019).

