## Extended data figures for "Mucosal Adenoviral-vectored Vaccine Boosting Durably Prevents XBB.1.16 Infection in Nonhuman Primates"

a

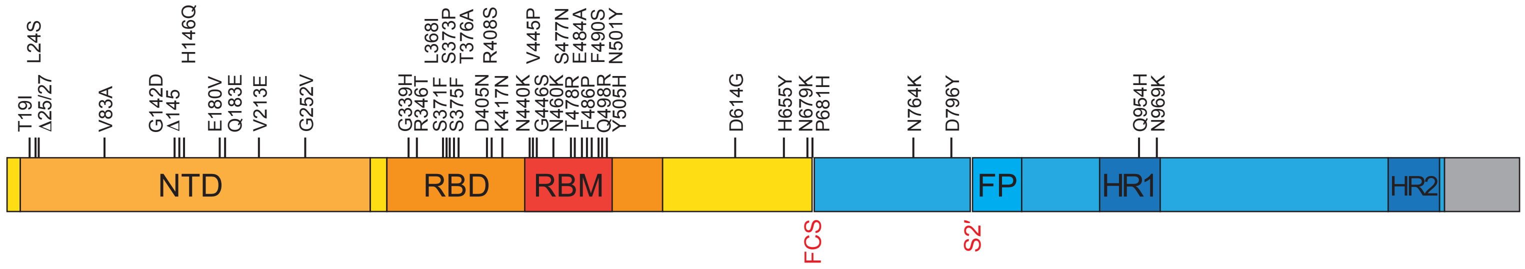


b

| Gene | Amino Acid Change |
|---|---|
| ORF1a | K47R |
| ORF1a | S135R |
| ORF1a | A307V |
| ORF1a | K680N |
| ORF1a | T842I |
| ORF1a | G1307S |
| ORF1a | L3027F |
| ORF1a | T3090I |
| ORF1a | L3201F |
| ORF1a | T3255I |
| ORF1a | P3395H |
| ORF1a | Δ3675/3677 |
| ORF1a | L3829F |
| ORF1b | P314L |
| ORF1b | G662S |
| ORF1b | S959P |
| ORF1b | M1197I |
| ORF1b | R1315C |
| ORF1b | I1566V |
| ORF1b | D1746Y |
| ORF1b | T2163I |
| ORF3a | T223I |
| E | T9I |
| E | T11A |
| M | Q19E |
| M | A63T |
| ORF6 | D61L |
| ORF8 | G8* |
| ORF8 | C83Y |
| N | P13L |
| N | Δ31/33 |
| N | R203K |
| N | G204R |
| N | S413R |

**Extended Data Figure 1**

**Extended Data Figure 1. XBB.1.16 sequence**

Challenge virus was titered and sequenced to verify presence of canonical XBB.1.16 amino acid substitutions in comparison to ancestral Wuhan-Hu-1 strain. Substitutions shown for (**a**) S gene and (**b**) whole genome. NTD: N-terminal domain. RBD: receptor binding domain. RBM: receptor binding motif. FP: fusion peptide. HR1: heptad repeat 1. HR2: heptad repeat 2. FCS: furin cleavage site. S2': S2' site. *: premature stop codon in Orf8 (common to XBB descendant strains).

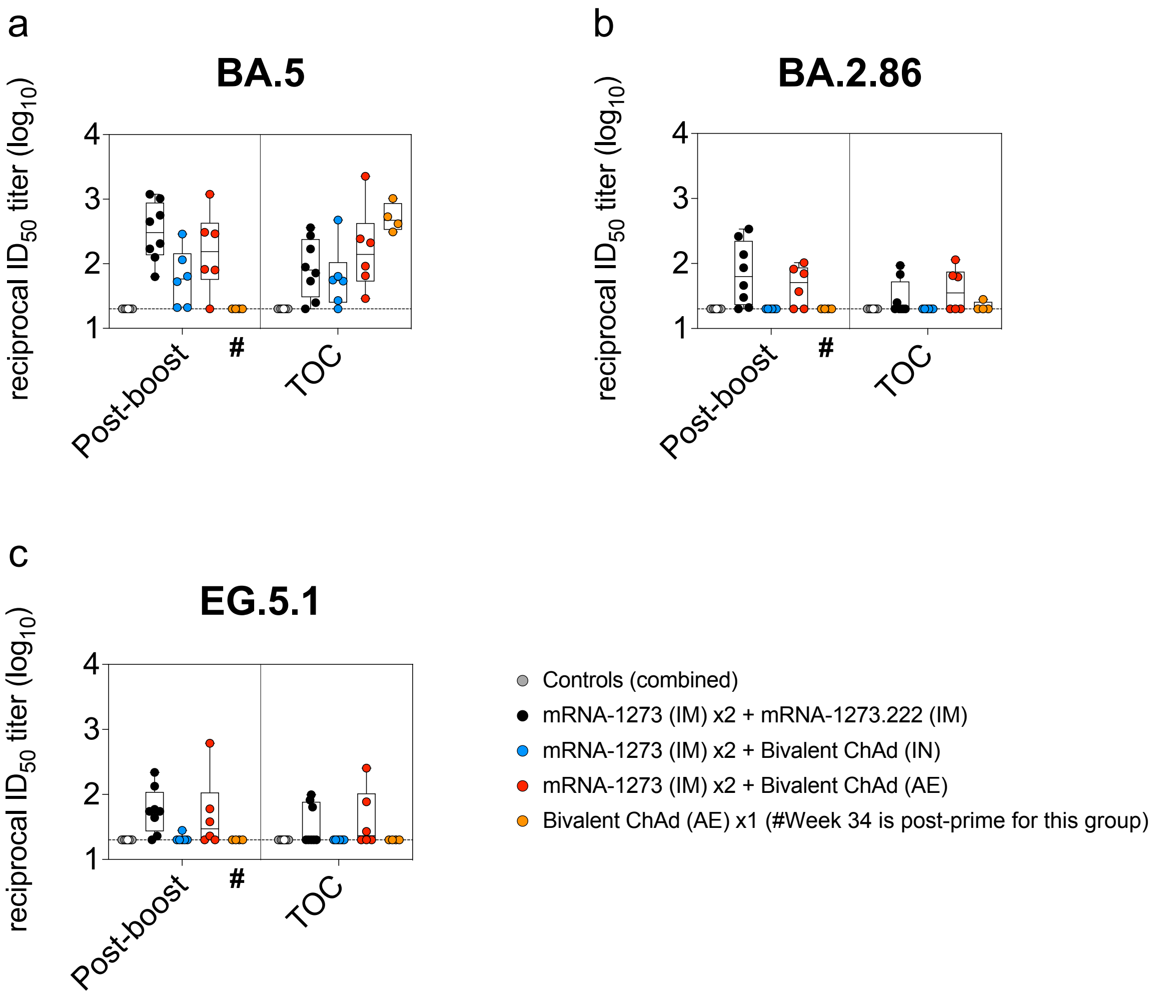


**Extended Data Figure 2. AE or IM boosting elicits durable serum neutralizing responses against emerging SARS-CoV-2 variants**

NHP (*n*=4-8 per group) were administered mRNA-1273 or control mRNA at weeks 0 and 4 and boosted at week 32 with the indicated vaccine. (**a-c**) Sera were collected post-boost (week 34) and at the time of challenge (week 48). Pseudovirus neutralizing responses measured against (**a**) BA.5, (**b**) BA.2.86 and (**c**) EG.5.1. Circles represent individual NHP. Boxes represent interquartile range with the median denoted by a horizontal line. Assay LOD represented as dotted line. # indicates that while sample collection for AE prime cohort (orange) occurred on week 34, week 34 was two weeks following the single AE prime rather than two weeks following a boost as in other groups.

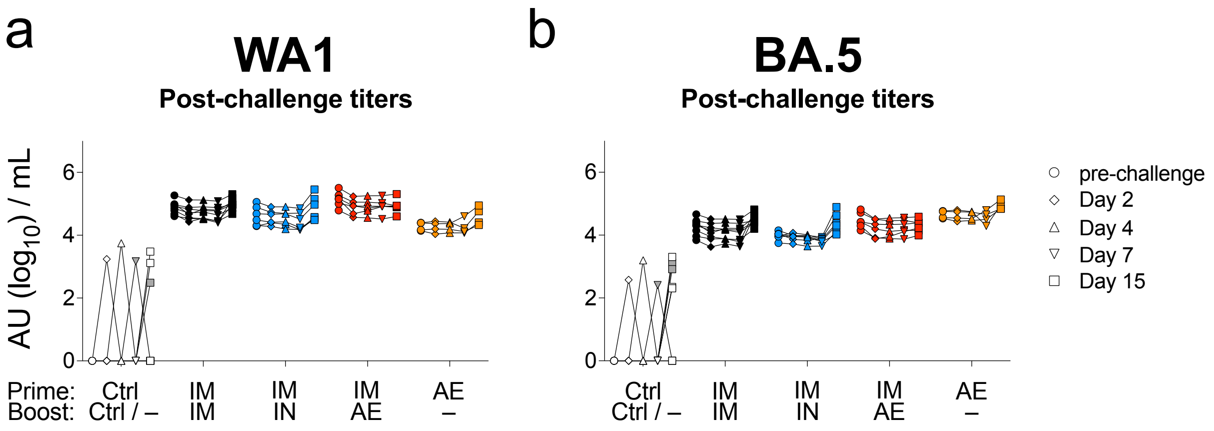


**Extended Data Figure 3. Post-challenge anamnestic responses in sera**

NHP (*n*=4-8 per group) were administered mRNA-1273 or control mRNA at weeks 0 and 4 and boosted at week 32 with the indicated vaccine. (**a-b**) Sera were collected pre-challenge (week 48) and post-challenge (days 2, 4, 7 and 15). (**a**) Anti-WA1 S and (**b**) anti-BA.5 S binding titers shown at indicated times. Symbols indicate AU / mL of individual NHP at indicated times. AU below a value of 1 were replaced with a value of 1.

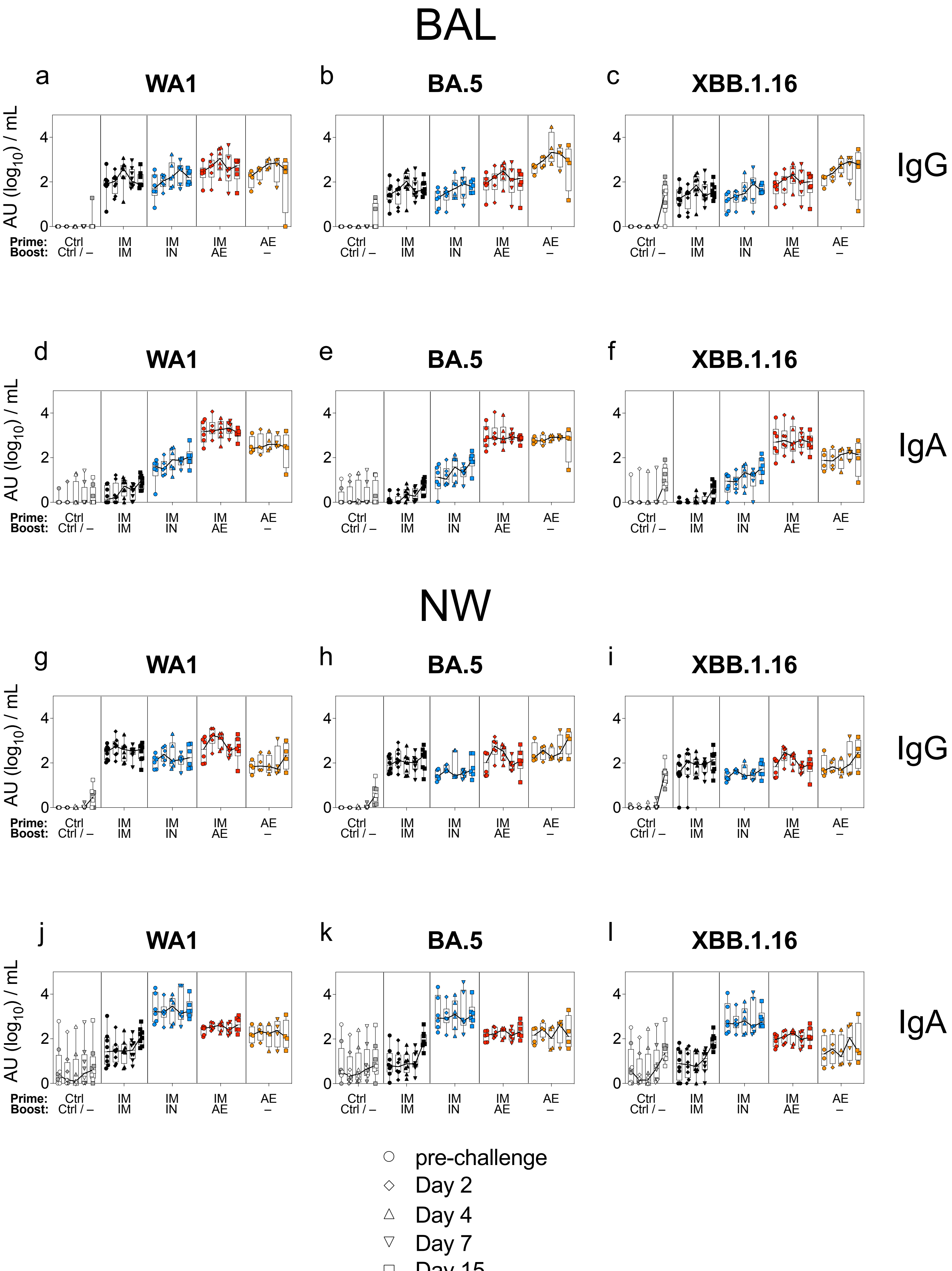


Extended Data Figure 4

**Extended Data Figure 4. XBB.1.16 challenge elicits rapid anamnestic antibody responses in the BAL of vaccinated NHP**

NHP (*n*=4-8 per group) were administered mRNA-1273 or control mRNA at weeks 0 and 4 and boosted at week 32 with the indicated vaccine. (**a-f**) BAL and (**g-l**) NW were collected pre-challenge (week 48) and on days 2, 4, 7 and 15 post-challenge. (**a-c**, **g-i**) IgG and (**d-f**, **j-l**) IgA binding titers measured using WA1, BA.5 or XBB.1.16 S as indicated. Symbols indicate individual NHP. Boxes represent interquartile range with median indicated by thin solid line. Thick solid lines connect median binding titers across timepoints. AU below a value of 1 were replaced with a value of 1.

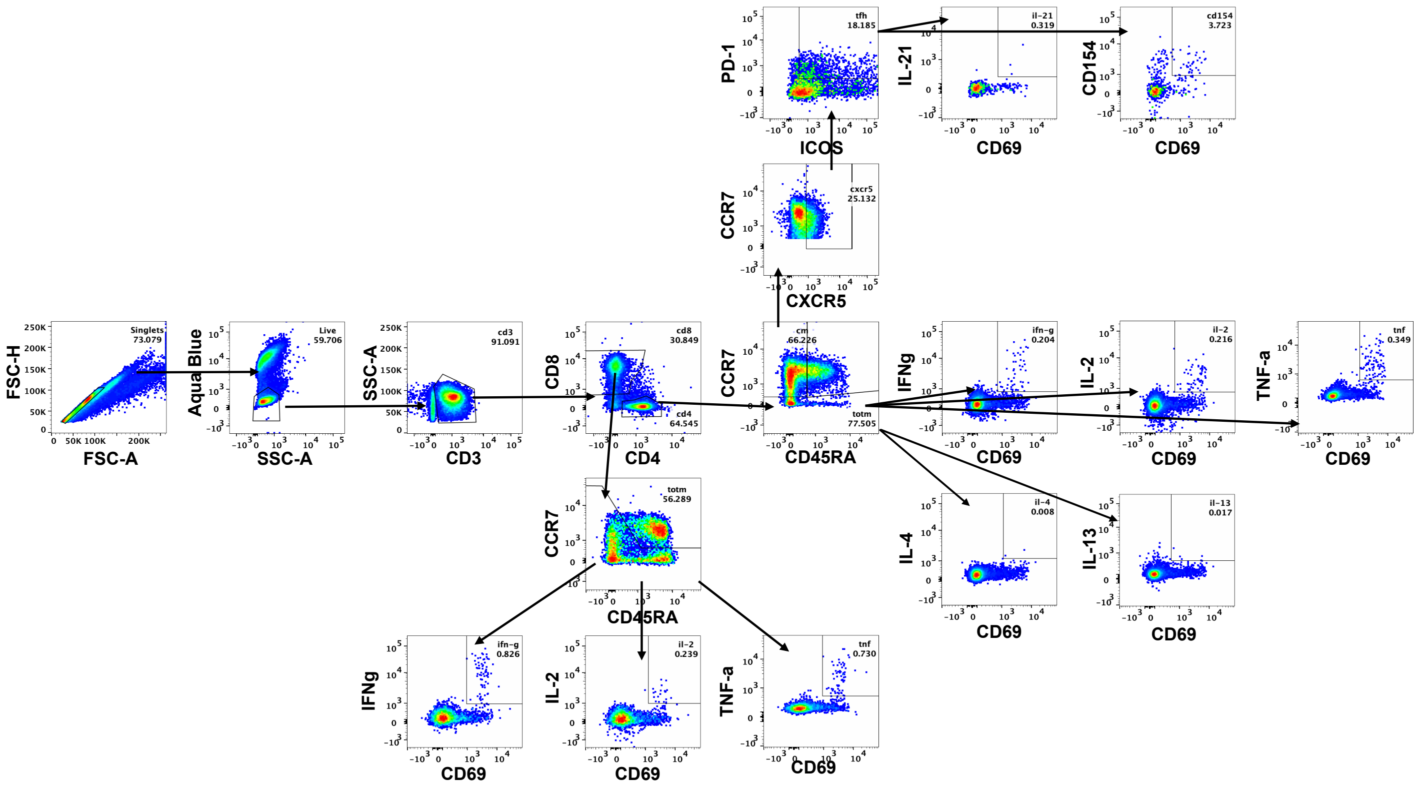


**Extended Data Figure 5. T cell gating strategy**

Representative flow cytometry plots showing gating strategy for ICS following S or N peptide stimulation. Cells were gated as singlets and live cells on forward and side scatter and a live/dead aqua blue stain. $CD3^+$ events were gated as $CD4^+$ or $CD8^+$ T cells. Total memory $CD8^+$ T cells were selected based on expression of CCR7 and CD45RA. SARS-CoV-2-specific memory $CD8^+$ T cells were gated according to co-expression of CD69 and IL-2, TNF-α or IFN-γ. The $CD4^+$ events were defined as naïve, total memory or central memory according to expression of CCR7 and CD45RA. $CD4^+$ cells with a $T_H1$ phenotype were defined as memory cells that co-expressed CD69 and IL-2, TNF-α or IFN-γ. $CD4^+$ cells with a $T_H2$ phenotype were defined as memory cells that co-expressed CD69 and IL-4 or IL-13. $T_{FH}$ cells were defined as central memory $CD4^+$ T cells that expressed CXCR5, ICOS and PD-1. $T_{FH}$ cells were further characterized as IL-$21^+$, $CD69^+$ or $CD40L^+$, $CD69^+$.

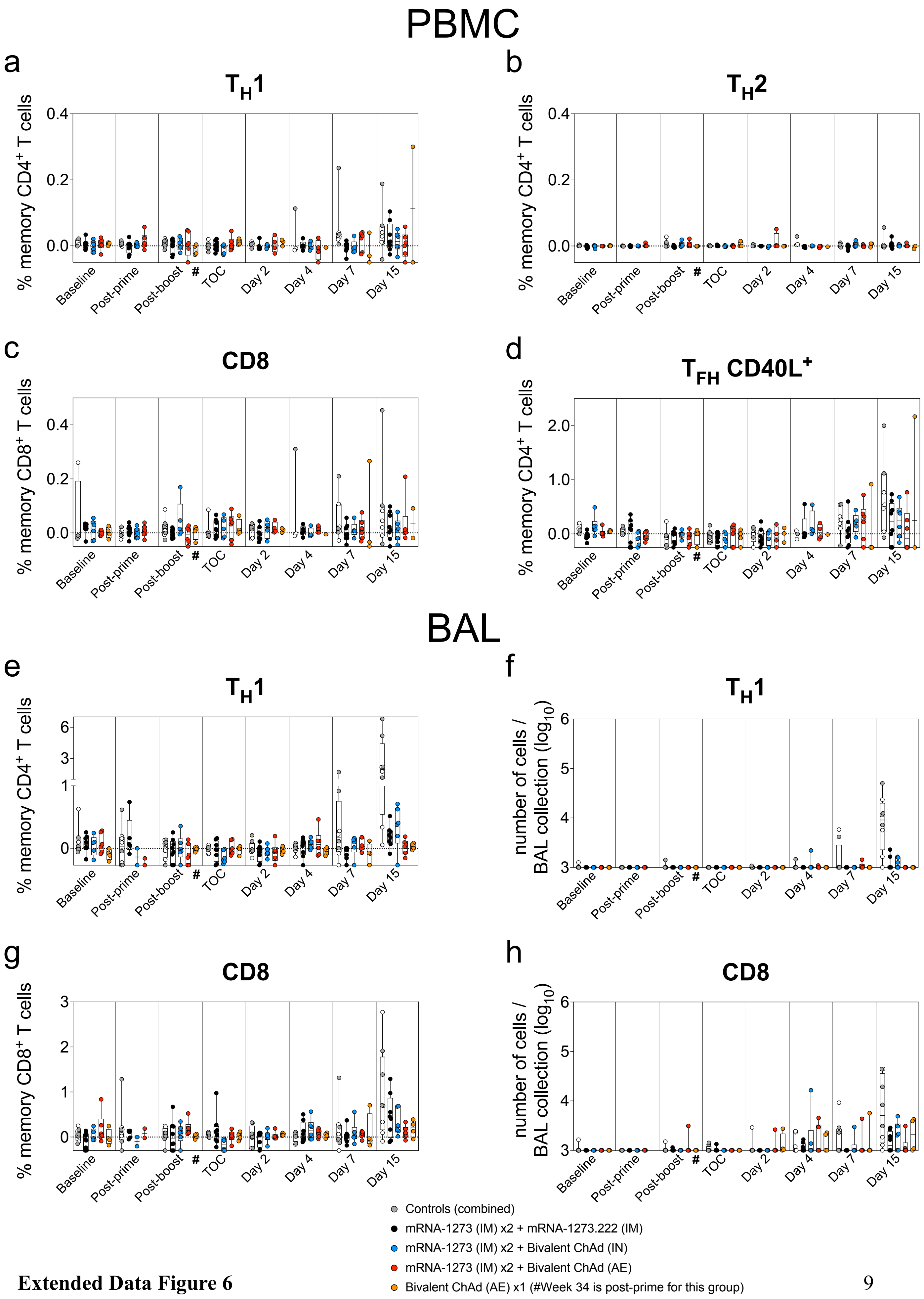


**Extended Data Figure 6**

**Extended Data Figure 6. T cell responses to N peptides in PBMC and BAL following IM or mucosal boosting**

(**a-d**) PBMC and (**e-h**) BAL fluid were collected pre-vaccination (baseline) and at weeks 6 (post-prime), 34 (post-boost) and 48 (time of challenge) as well as on days 2, 4, 7 and 15 post-challenge. Lymphocytes were stimulated with SARS-CoV-2 N peptides (WA1) and then measured by intracellular staining. (**a-b, e**) Percentage of memory $CD4^+$ T cells with (**a, e**) $T_H1$ markers (IL-2, TNF-$\alpha$ or IFN-$\gamma$) or (**b**) $T_H2$ markers (IL-4 or IL-13) following stimulation. (**c, g**) Percentage of memory $CD8^+$ T cells expressing IL-2, TNF-$\alpha$ or IFN-$\gamma$ following stimulation. (**d**) Percentage of $T_{FH}$ cells that express CD40L following stimulation. Break in y-axis in **e** indicates a change in scale without a break in the range depicted. Dotted lines set at 0%.

Absolute number of N-reactive (**f**) $T_H1$ $CD4^+$ or (**h**) $CD8^+$ T cells in the BAL are also depicted. Counts below a value of 1 (due to background subtraction) were replaced with a value of 1 for data in **f** and **h**. Circles in **a-h** indicate individual NHP. Boxes represent interquartile range with the median denoted by a horizontal line. Reported values may be negative due to background subtraction and may extend below the range of the y-axis. # indicates that while sample collection for AE prime cohort (orange) occurred on week 34, week 34 was two weeks following the single AE prime rather than two weeks following a boost as in all other groups.
